# A biatrial digital twin integrating electrophysiology, mechanics, and circulation: from physiology to atrial fibrillation

**DOI:** 10.64898/2026.03.12.711092

**Authors:** Sergi Picó Cabiró, Alberto Zingaro, Violeta Puche García, Dimitrios Lialios, Mariano Vázquez, Blas Echebarria, Maite Izquierdo, Francesc Carreras, Javier Saiz, Eva Casoni

## Abstract

Atrial electromechanics plays a key role in cardiac function by regulating ventricular filling and global hemodynamics, yet remains challenging to model consistently across scales. In this work, a multiscale atrial digital twin for simulations of normal and pathological atrial function is presented, formulated as an electromechanical framework for biatrial simulations that couples three-dimensional atrial electrophysiology and mechanics with a closed-loop zero-dimensional circulatory model. The framework is calibrated on a patient-specific biatrial anatomy to reproduce physiological regional activation times, atrial volumes, ejection fractions, and pressure–volume loop characteristics. The simulations capture all atrial functional phases throughout the cardiac cycle, including realistic figure-eight pressure–volume loops, an aspect hard to achieve in computational studies. A systematic sensitivity analysis quantifies the influence of active contraction, passive stiffness, boundary conditions, and circulatory parameters on atrial function. Finally, application to a pathological scenario through induced persistent atrial fibrillation demonstrates how electrophysiological remodelling propagates across scales, leading to loss of effective atrial contraction, altered atrioventricular flow patterns, and a clinically relevant reduction in cardiac output. Overall, this multiphysics and multiscale framework provides a robust platform to investigate how atrial electrical alterations drive mechanical and hemodynamic alterations in both healthy and pathological conditions.

## 1. Introduction

The atria are not simply passive conduits for blood flow but dynamic heart chambers that actively contribute to ventricular filling and overall cardiac output (CO) [1]. In pathological conditions such as atrial fibrillation (AF), the most prevalent sustained cardiac arrhythmia, disruption of atrial electrical and mechanical activity leads to impaired hemodynamics and elevated risk of complications [2]. According to recent global burden data, AF affected an estimated 52.5 million individuals worldwide in 2021 [3]. Standard treatments such as antiarrhythmic drugs and catheter ablation aim to restore sinus rhythm and anticoagulation is mandatory to prevent stroke. However, recurrence remains high; up to 40–50% of patients relapse within five years of initially successful ablation [4]. These limitations highlight the need for more personalized and predictive strategies in AF management, with *in silico* models offering a valuable tool to address these challenges. Computational models of the human atria have been developed to investigate atrial function under physiological and pathological conditions, with varying focuses and methodological approaches [5, 6, 7, 8, 9, 10, 11, 12, 13, 14, 15, 16].

The majority of these approaches are built on electrophysiological models, which simulate electrical activation and propagation in cardiac tissue, thereby enabling the investigation of the mechanisms underlying arrhythmia initiation and maintenance [17, 7, 18, 6, 19, 20, 21, 22, 23, 24, 5]. These frameworks have provided insights into fibrosis, conduction heterogeneity, and ablation strategies [6, 19, 25], and have also been applied to virtual drug testing [26, 27, 28, 23]. Computational fluid dynamics (CFD) models have, in parallel, been used to study atrial hemodynamics and thrombus risk, with particular focus on the left atrial appendage (LAA) [8, 29, 9, 30, 31]. These works highlight how altered flow patterns in AF promote stasis, and how interventions such as LAA occlusion can be assessed *in silico* [32].

Electromechanical models integrate electrophysiology with active force generation and tissue mechanics, allowing simulation of how activation patterns drive atrial motion. The first 3D biatrial electromechanical model was developed by Adeniran *et al*. [10], incorporating heterogeneous cellular properties, anisotropic conduction, and AF-induced electrical remodelling, and showing that these alterations markedly reduce calcium transients, active force, and ejection fraction (EF). Land and Niederer [11] incorporated atrial-specific contraction dynamics in a four-chamber framework to examine atrial–ventricular interaction and post-cardioversion recovery. Patient-specific geometries have enabled stress and strain analysis, as in Augustin *et al*. [33], who quantified the influence of wall thickness and curvature on left atrial wall stress, while Gerach *et al*. and Fedele *et al*. [34, 35] examined whole-heart electromechanical simulations integrating atrial and ventricular mechanics with closed-loop 0D circulation models. Furthermore, Strocchi *et al*. [36] developed a four-chamber electromechanical computational model to identify parameters most affecting clinically relevant biomarkers. By linking electrical activation, mechanical contraction, and hemodynamic loading, electromechanical frameworks can address major limitations of purely electrophysiological or CFD models, such as their inability to capture the sequence from conduction disturbances through mechanical dysfunction to the resulting hemodynamic alterations within the atria. However, significant challenges remain unsolved. Several studies have emphasized that validation is limited by the lack of clinical datasets containing mechanical biomarkers [10, 11, 33]. Others highlight that reproducing realistic atrial pressure–volume loops is challenging [35], and some models fail to capture them accurately, producing unrealistic loop morphologies [34, 37, 38]. Finally, key parameters, including those describing passive mechanics and atrioventricular boundary conditions, remain uncertain, as highlighted in prior studies [33, 36, 34].

In this paper, a 3D electromechanical–0D hemodynamic biatrial digital twin model is presented. The framework integrates region-specific atrial electrophysiology, region-dependent diffusion values calibrated from activation times to reproduce physiological activation times. Atrial-specific active contraction dynamics are incorporated within a solid mechanics formulation and coupled to a 0D circulatory model. It captures all phases of atrial function and produces mechanical biomarkers in close agreement with values reported in the literature, including realistic figure-eight pressure–volume loops. In addition, a comprehensive sensitivity analysis is performed, in which parameters are systematically varied across different model components: electrophysiological diffusion coefficients, elastance parameters of the circulation model, maximal active tension at rest length, and passive mechanical properties of atrial tissue. Variations in model parameters are evaluated in terms of their impact on clinically relevant biomarkers [39, 40], including atrial EFs and chamber volumes over the cardiac cycle. The framework is further tested in a pathological scenario by reproducing AF in persistent conditions, combining an electrical remodelling scheme with an S1–S2 stimulation protocol [41, 42]. These results demonstrate the versatility of the framework for investigating disease-induced alterations in atrial function.

The main contribution of this work with respect to previous studies is the combination of capabilities this 3D electromechanical–0D hemodynamic model provides: quantitative reproduction of regional activation times, atrial volumes, and pressure–volume loop morphology within a fully coupled electromechanical-circulatory setting; simulation of persistent AF within a three-dimensional electromechanical atrial model integrated with a 0D closed-loop circulation; and a systematic sensitivity analysis of atrial mechanical and hemodynamic parameters. Together, these capabilities advance prior electromechanical models by unifying biatrial electrophysiology, solid mechanics, and circulation, enabling calibration against clinical biomarkers and capturing the direct electromechanical and hemodynamic consequences of AF.

## 2. Methods

A 3D biatrial electromechanical model coupled to a 0D closed-loop circulation was developed from patient-specific atrial anatomy. The framework incorporates regional cellular heterogeneities and atrial-specific mechanical properties to reproduce physiologically realistic hemodynamic conditions. Starting from computed tomography data, the atrial geometry was reconstructed through regional segmentation and surface processing. The resulting surface was volumetrically meshed to define the computational domain, and myocardial fiber orientations were subsequently assigned. Simulations were performed on a high-performance computing (HPC) platform. Quantitative metrics, including electrical activation times, atrial volume dynamics, and EFs, were extracted for calibration and analysis.

### 2.1. Anatomical biatrial model

#### 2.1.1. Imaging data acquisition

The anatomical geometry was obtained from a contrast-enhanced computed tomography scan acquired at Hospital Universitari i Politècnic La Fe (Valencia, Spain). The dataset corresponded to a 56-year-old male patient with a history of hypertension and recurrent atrial arrhythmias, including documented episodes of typical atrial flutter. Image acquisition was performed during ventricular diastole to reduce cardiac motion and improve delineation of atrial anatomy.

#### 2.1.2. Anatomical biatrial reconstruction

A patient-specific three-dimensional biatrial geometry was reconstructed by segmenting the atrial anatomical structures using Seg3D ^1^ (Fig 1). Automatic segmentation was complemented by manual corrections to ensure anatomical consistency. The resulting surface model was smoothed using filters implemented in ParaView ^2^ and Blender ^3^.

**Figure 1:**
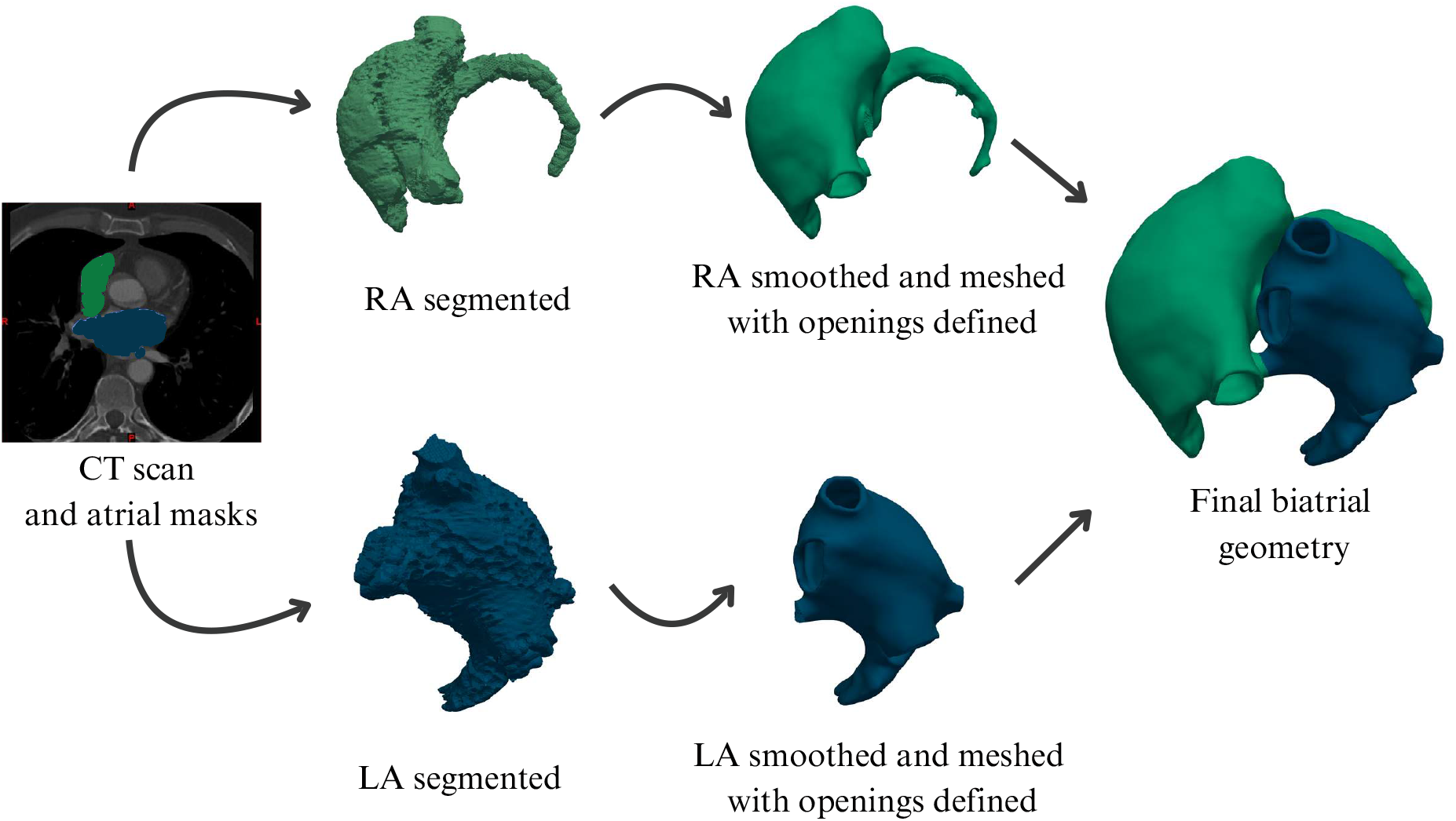
Anatomical reconstruction. Computed tomography based atrial segmentation, surface processing, and assembly of the final biatrial geometry.

Atrioventricular valve openings and atrial connections were then identified and incorporated into the surface model. A region-dependent atrial wall thickness was explicitly imposed during geometric construction of the volumetric anatomy, based on anatomical and imaging evidence demonstrating marked heterogeneity across atrial regions, particularly in the right atrium [43, 44, 45]. Incorporating this imposed heterogeneity, rather than assuming a homogeneous thickness as in earlier atrial models [46, 47, 48, 49], improves anatomical fidelity and supports more physiologically grounded simulations [50, 51, 52].

To represent anatomical and functional heterogeneity, the atrial geometry was subdivided into 48 distinct anatomical regions, following regional classification [53]. Region boundaries were defined using a semi-automatic algorithm based on anatomical landmarks and surface topology [54]. This regionalization enabled region-specific assignment of fiber orientation, cellular electrophysiology, and tissue conductivities.

Myocardial fiber orientation was prescribed using a region-based, rule-driven approach. For each anatomical region, a principal region-defining direction vector was defined following detailed atrial anatomical descriptions from published atlases, in particular Sánchez-Quintana et al. [55]. This vector was used as a regional anatomical reference to compute local fiber directions. In atrial wall regions and the Bachmann bundle, fibers were obtained by projecting the region-defining vector onto the local surface tangent plane, resulting in longitudinal fiber orientations aligned with the dominant anatomical direction. In contrast, in appendages, valves, and orifices, a transverse arrangement was prescribed by computing fiber directions as the cross product between the region-defining vector and the local surface normal, yielding tangential, circumferential orientations consistent with atrial anatomy [53]. The resulting fiber architecture is illustrated in Fig 2.

**Figure 2:**
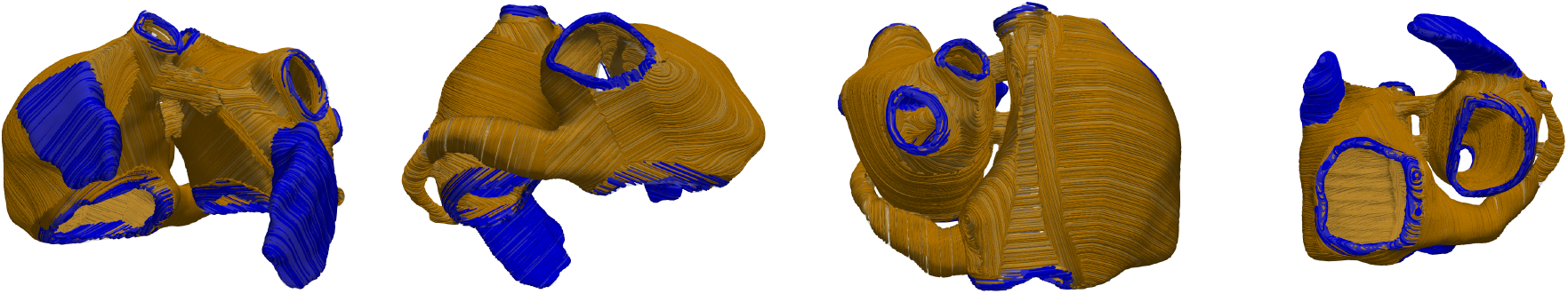
Rule-based atrial myocardial fiber architecture. Fiber orientations across the left and right atria. Colours indicate regions where fibers are prescribed using different orientation rules (longitudinal wall/bundle alignment versus circumferential appendage and hole alignment).

#### 2.1.3. Mesh generation

Volumetric meshes were generated from the reconstructed atrial geometry to define the computational discretization. To balance numerical accuracy and computational cost, two meshes were employed: a fine mesh for electrophysiology simulations and a coarser mesh for solid mechanics (Fig 3).

**Figure 3:**
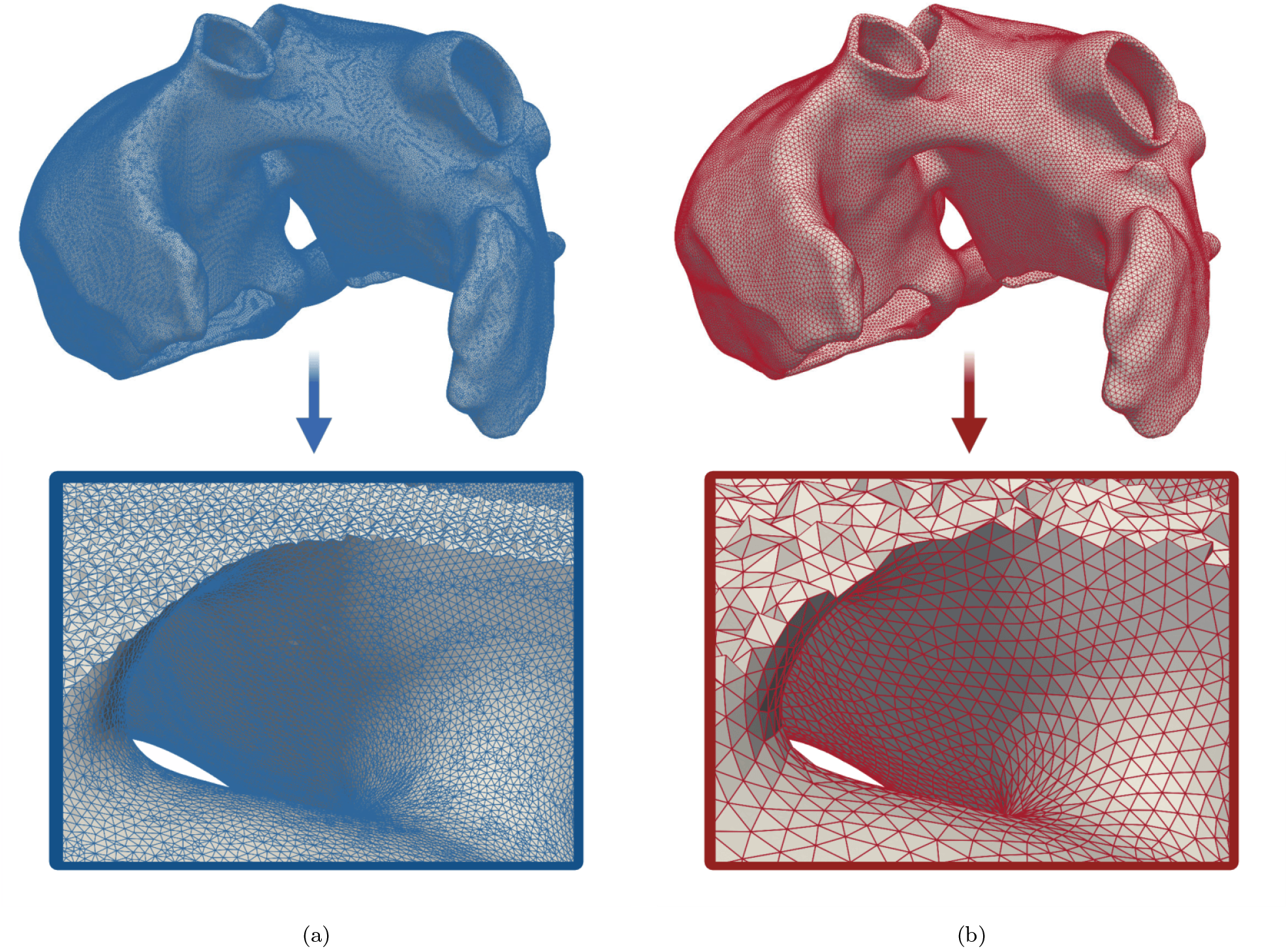
Meshes used in the electromechanical simulation. (a) Electrophysiology mesh. (b) Solid mechanics mesh. Zoomed views highlight the difference in spatial resolution between the fine mesh used for electrophysiology and the coarser mesh used for mechanics.

Electrical wavefront propagation exhibits steep spatial gradients requiring high spatial resolution, whereas mechanical deformation evolves more smoothly and can be accurately represented on a coarser discretization. This two-mesh strategy was implemented using a multi-instance framework [56].

The electrophysiology mesh was generated using Quartet ^4^, which employs an isosurface-stuffing algorithm [57] to produce regular volumetric elements. The solid mechanics mesh was obtained by remeshing and coarsening the electrophysiology discretization in ANSA^5^. Mesh characteristics are summarized in Table 1. The average edge length was 0.267 ± 0.038 mm for electrophysiology and 0.799 ± 0.148 mm for mechanics. These values are consistent with spatial resolutions reported in atrial and whole-heart modeling studies [58, 59].

**Table 1:** Mesh resolution and tetrahedral quality metrics for the electrophysiology and solid mechanics discretizations.

|  | **Electrophysiology mesh** | **Solid mechanics mesh** |
| --- | --- | --- |
| Number of tetrahedra | 16,730,066 | 718,882 |
| Average edge length (mm) | $0.267 \pm 0.038$ | $0.799 \pm 0.148$ |
| Average quality | 0.945 | 0.873 |
| Minimum quality | 0.316 | 0.469 |

Mesh quality was evaluated using a normalized tetrahedral quality metric ranging from 0 (degenerate element) to 1 (equilateral tetrahedron) [60, 61]. The electrophysiology mesh exhibited a mean quality of 0.945 (minimum 0.316), reflecting the regularity characteristic of isosurface-stuffing–based meshing. The solid mechanics mesh exhibited a mean quality of 0.873 (minimum 0.469), slightly lower due to coarsening. Although minimum values indicate moderate distortion in some elements, all elements remained well within acceptable ranges for stable numerical simulation [60].

### 2.2. The 3D electromechanical–0D hemodynamic model

Fig 4 presents a general overview of the cardiac modelling framework. The model is multiphysics, as it integrates electrical propagation, active and passive tissue mechanics, and hemodynamics within a single formulation. It is also multiscale, in both physiological and geometrical perspectives. The framework spans multiple biological scales, ranging from subcellular ionic currents and calcium dynamics, through cellular excitation-contraction coupling and tissue-level electrical propagation, to active force generation and whole-organ pump function. From a geometrical perspective, it combines 3D finite-element representations of atrial electrophysiology and solid mechanics with a zero-dimensional description of the circulatory system [62]. The model is strongly coupled, since each physical description interacts consistently with the others with feedback mechanisms within a unified modeling framework. EP drives intracellular calcium transients, and these transients regulate active force generation. Mechanical contraction then determines atrial deformation and associated pressure–volume relations. These pressures and volumes are consistently linked to the 0D circulatory model, which represents both ventricular dynamics and the systemic and pulmonary circulations. As a result, the model captures not only atrial electromechanical behaviour in three dimensions, but also its functional integration within the global circulatory system.

**Figure 4:**
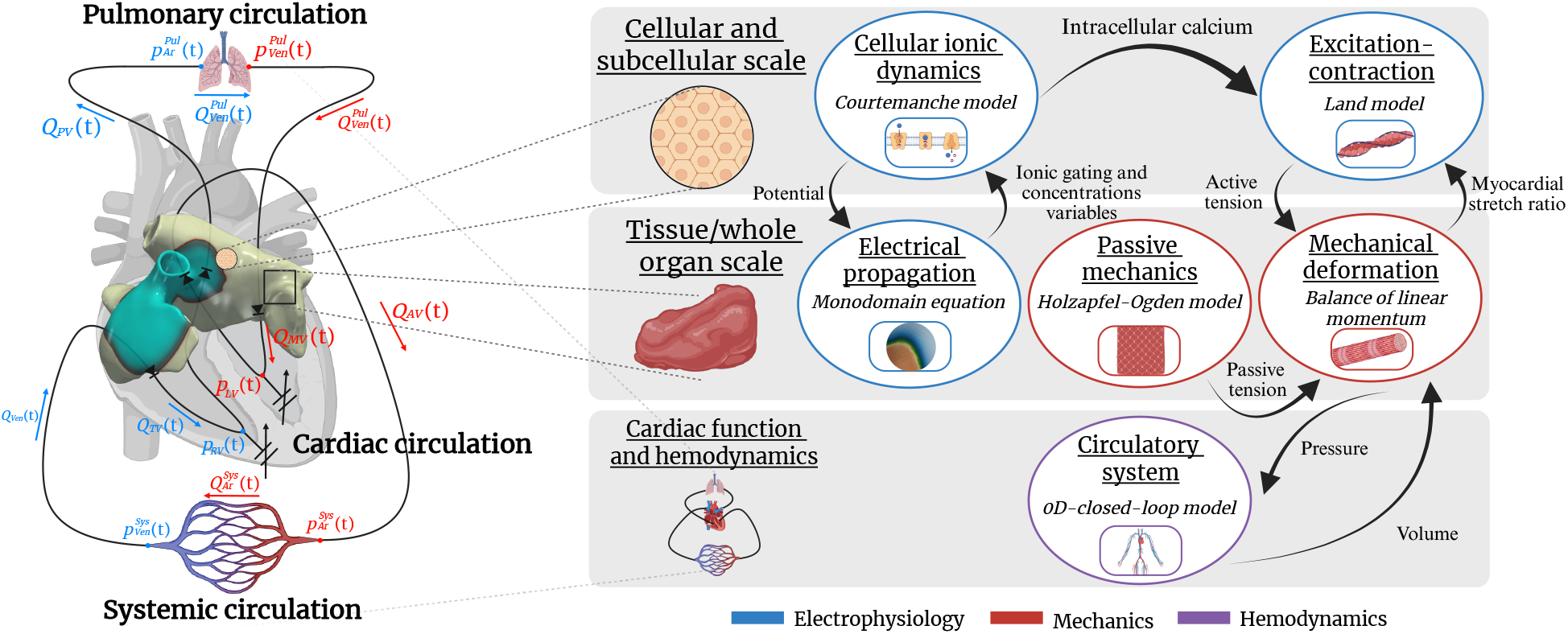
Multiscale and multiphysics model of the atria.

Most constitutive parameters in the different submodels are taken from previously published works: cellular electrophysiology follows the human atrial model of Courtemanche– Ramirez–Nattel (CRN) [63], passive atrial mechanics followed the atrial parameterization proposed by Monaci *et al*. [64], active tension generation is based on the excitation-contraction formulation of Land *et al*. [65, 11], and the 0D closed-loop circulation is derived from models stablished the models of Blanco *et al*. [66] and Regazzoni *et al*. [67]. However, a subset of parameters is calibrated within physiologically plausible ranges to ensure that the coupled 3D-0D model reproduces realistic regional atrial activation patterns, chamber volumes, and EFs.

#### 2.2.1. Electrophysiology

Electrical activation at the tissue level is modeled with the monodomain equation, which describes the spread of the transmembrane potential through cardiac tissue. This equation is expressed as [68]:

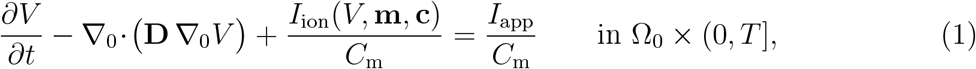

where Ω_0_ is the reference atrial domain at *t* = 0, *T* the final time, *V* the transmembrane potential, *C*_*m*_ the membrane capacitance per unit area, **D** the diffusion tensor, *I*_app_ an external stimulus, applied in this work at the sinoatrial node (SAN), and ∇_0_ the nabla differ ential operator with respect to the reference configuration. The total ionic current *I*_ion_ = ∑_*i*_ *I*_*i*_, together with the state variables **m** (gating variables) and **c** (ionic concentrations) governing the local action potential, were defined according to the selected cellular ionic model.

Tissue-level electrical conduction was represented through an orthotropic diffusion tensor aligned with the local fiber coordinate system,

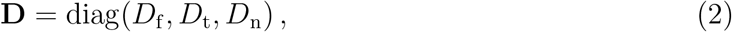

with longitudinal (*D*_f_), transverse (*D*_t_), and normal (*D*_n_) components. Transverse and normal diffusion coefficients were expressed using a single anisotropy ratio *r* ∈ (0, 1] [53],

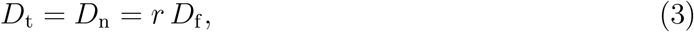

thereby reducing the number of independent diffusion parameters. The diffusion coefficients were subsequently calibrated to reproduce physiological activation times.

##### Cellular ionic model

Cellular atrial electrophysiology was represented using the CRN [63] model, a widely used human atrial formulation incorporating detailed sodium, potassium, and calcium fluxes together with sarcoplasmic reticulum cycling. This model has been employed extensively in tissue-and organ-scale simulations of atrial electrophysiology [24, 54, 34] and in studies of arrhythmogenic mechanisms [5, 69, 20]. The acetylcholine-activated potassium current *I*_KACh_ was included [70], with [ACh] = 0.005 *µ*M [5].

Electrophysiological heterogeneity was represented using two distinct sets of anatomical regions: one associated with tissue-level conduction properties in the electrical propagation model, and another associated with cellular-level ionic properties in the ionic model. These regional subdivisions were defined independently, as conduction heterogeneity and ionic heterogeneity arise from different underlying mechanisms and therefore do not necessarily coincide anatomically [53]. The two sets of regions employed in this work are illustrated in Fig 5.

**Figure 5:**
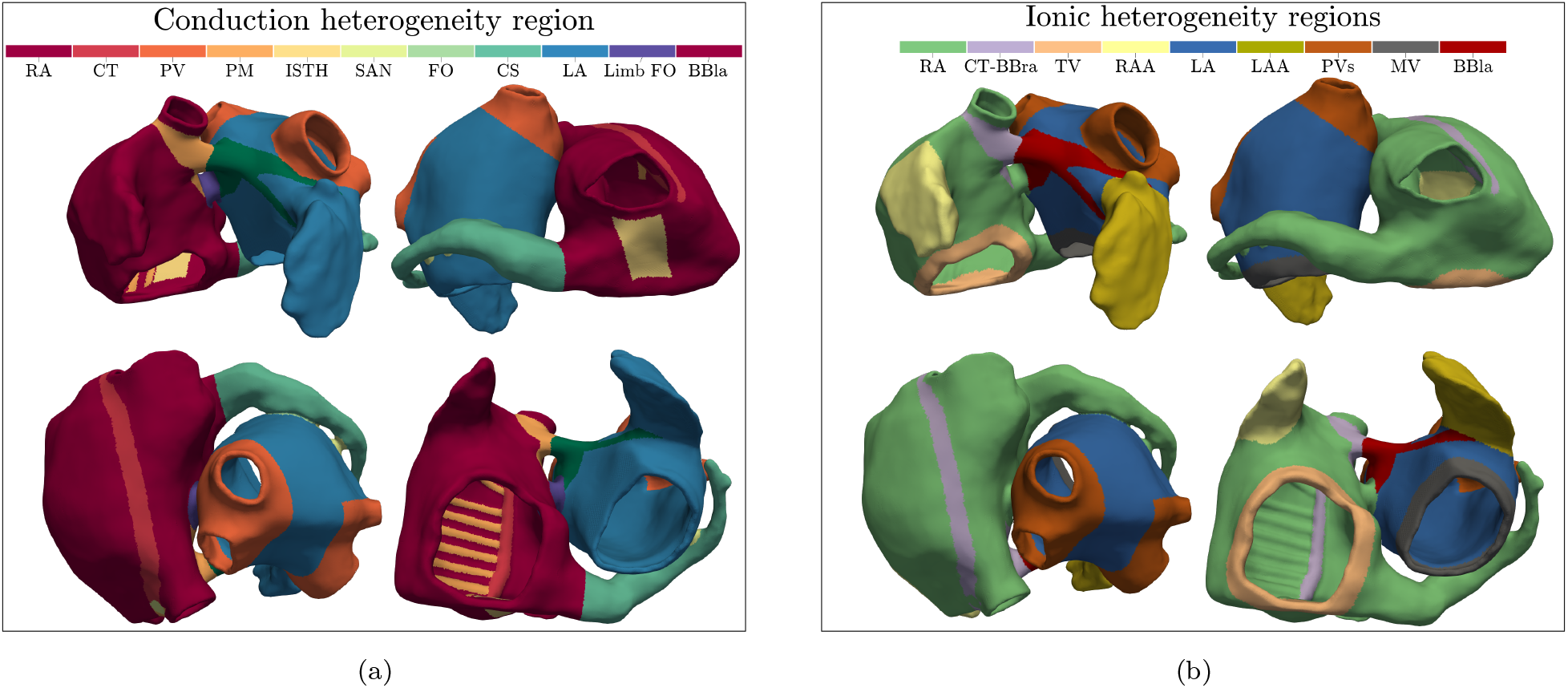
Electrophysiological regional heterogeneity used in the model. (a) Regions used to prescribe conduction properties in the electrical propagation model. (b) Regions used to prescribe ionic heterogeneity in the cellular ionic model.

##### Regional electrical conduction heterogeneity

Conduction velocity varies across atrial structures [71, 72, 73]. To account for this, the atria are partitioned into eleven anatomical regions (Fig 5a): right atrium (RA), crista terminalis (CT), pulmonary veins (PVs), pectinate muscles (PMs), cavotricuspid isthmus (ISTH), SAN, fossa ovalis (FO), coronary sinus (CS), left atrium (LA), limbus of the fossa ovalis (FO limb), and Bachmann’s bundle of the left atrium (BBla). Each region is characterized by an orthotropic diffusion tensor aligned with the local fiber basis.

The anisotropy ratio *r* appearing in Eq. (3) is prescribed for each region based on prior modeling investigations of atrial electrical conduction anisotropy [53], which derived these quantities based on experimental observations [74, 75, 76, 48]. The adopted values are summarized in Table 2.

**Table 2:**
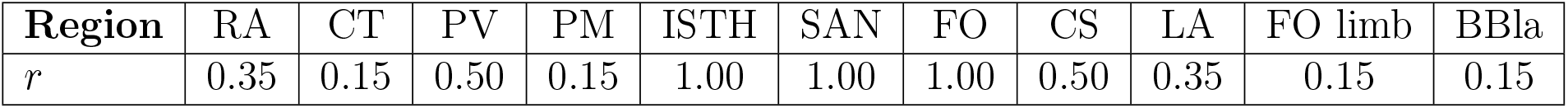
Regional anisotropy ratio *r* used in Eq (3). Values were adopted from prior atrial conduction modeling studies [53]. Regions with *r* = 1.00 correspond to isotropic conduction.

| Region | RA | CT | PV | PM | ISTH | SAN | FO | CS | LA | FO limb | BBla |
| --- | --- | --- | --- | --- | --- | --- | --- | --- | --- | --- | --- |
| $r$ | 0.35 | 0.15 | 0.50 | 0.15 | 1.00 | 1.00 | 1.00 | 0.50 | 0.35 | 0.15 | 0.15 |

##### Regional ionic heterogeneity

To capture experimentally observed ionic heterogeneity across the atria, selected maximal conductances were scaled in nine anatomical regions (Fig 5b): RA, CT–Bachmann’s bundle of the right atrium (BBra), tricuspid valve (TV), right atrial appendage (RAA), LA, LAA, PVs, mitral valve (MV), and BBla. The scaling factors relative to the baseline RA region of the CRN model are provided in Table 3, following previous studies [77, 5].

**Table 3:** Scaling factors for selected ionic conductances in different atrial regions relative to the CRN baseline right atrial model. Values were adopted from previous studies [77, 5].

| Region | RA | CT–BBra | TV | RAA | LA | LAA | PVs | MV | BBla |
| --- | --- | --- | --- | --- | --- | --- | --- | --- | --- |
| $g_{K1}$ | 1 | 1 | 1 | 1 | 1 | 1 | 0.9 | 1 | 1 |
| $g_{to}$ | 1 | 1 | 1 | 0.68 | 1 | 1 | 0.9 | 1 | 1 |
| $g_{Kr}$ | 1 | 1 | 1.3 | 1.1 | 2.0 | 2.2 | 2.5 | 2.6 | 1.6 |
| $g_{Ks}$ | 1 | 1 | 1 | 1 | 1 | 1 | 1.9 | 1 | 1 |
| $g_{CaL}$ | 1 | 1.67 | 0.8 | 0.9 | 0.9 | 0.9 | 0.8 | 0.8 | 1.67 |

#### 2.2.2. Solid mechanics

Atrial deformation was governed by the balance of linear momentum and mass conservation which, in the reference configuration, read as [68]:

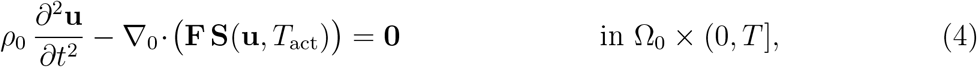

where **u** denotes the displacement field, *ρ*_0_ the reference density, **F** = **I**+ ∇_0_**u** the deformation gradient, **C** = **F**^⊤^**F** the right Cauchy–Green tensor, **S** the second Piola–Kirchhoff stress tensor and *T*_act_ the active tension. The second Piola–Kirchhoff stress tensor was decomposed into passive and active contributions:

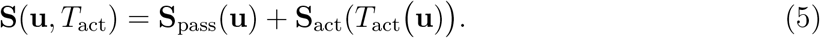

##### Passive mechanics

The passive mechanical behaviour of the tissue was represented using the transversely isotropic Holzapfel–Ogden hyperelastic law [78], with a volumetric penalty term to enforce quasi-incompressibility. Let *I*_1_ = tr**C** and *I*_4_ = **f**_0_ *·* **Cf**_0_ denote the invariants associated with the fiber direction **f**_0_. The strain–energy density function was defined as:

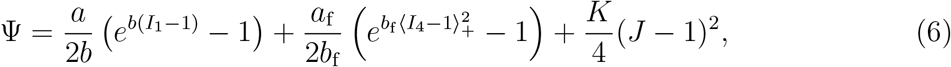

with ⟨*·*⟩_+_ = max(*·*, 0). The passive component of the second Piola–Kirchhoff stress was obtained as

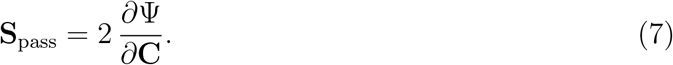

The coefficients *a* and *a*_f_ are stress-like parameters defining the baseline stiffness of the extracellular matrix and fiber families, whereas *b* and *b*_f_ are dimensionless stiffening parameters governing the exponential response. Their values correspond to the atrial adaptation of ventricular material properties described above and are reported in Table S1.

The bulk modulus *K* penalized deviations from incompressibility. A value of *K* = 1218.5 kPa was used, which was several orders of magnitude larger than the deviatoric stiffness parameters (*a, a*_f_), ensuring nearly incompressible behaviour consistent with the high water content of atrial myocardium. This formulation is consistent with cardiac mechanics models that treat myocardium as nearly incompressible using volumetric penalty terms [79, 10, 33]. The resulting myocardial volume variation remained small (∼1.8%), confirming near-incompressible behaviour without imposing strict isochoric constraints.

##### Active tension generation model

Active tension was computed using the excitation–contraction model of Land *et al*. [65], derived from human cardiomyocyte data and describing crossbridge dynamics through internal variables driven by intracellular calcium [Ca^2+^]_*i*_, fiber stretch *λ*_f_, and stretch rate 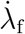.

The model introduced internal variables representing the fractions of blocked binding sites (*B*), weakly bound (*W*) and strongly bound (*S*) crossbridges, the mean distortion variables (*ζ*_*w*_, *ζ*_*s*_), and the fraction of calcium-bound troponin C (CaTRPN), collected in the state vector **q** = (*B, W, S, ζ*_*s*_, *ζ*_*w*_, CaTRPN), whose evolution obeys

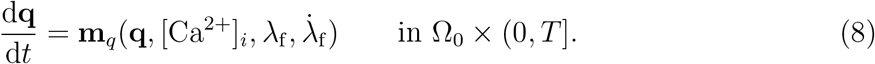

Active tension was defined as

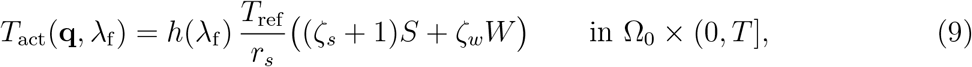

where *T*_ref_ denotes the reference maximal tension, *r*_*s*_ the steady-state fraction of strongly bound crossbridges, and *h*(*λ*_f_) the length-dependent modulation function. The active second Piola–Kirchhoff stress contribution was written as

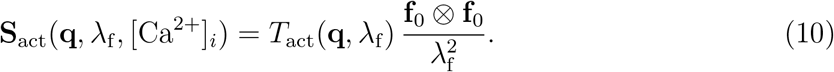

Although originally formulated for ventricular myocytes, the atrial reparameterization proposed by Land and Niederer [11] was adopted, adjusting kinetic parameters to reproduce atrial contractile properties. The reference tension *T*_ref_ was calibrated within the electromechanical framework to reproduce physiologically realistic atrial mechanics. Parameter values and sources are provided in S2 Table.

##### Boundary conditions

Robin and Neumann type boundary conditions were imposed to the mechanical model on distinct regions of the reference boundary *∂*Ω^0^, namely

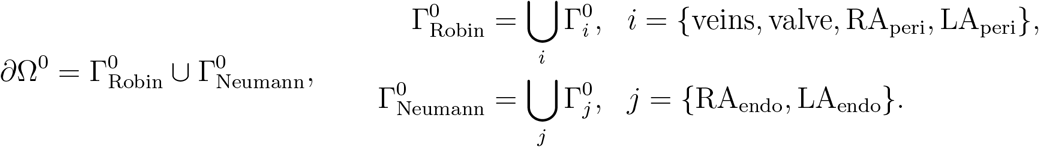

The corresponding boundary conditions were expressed as

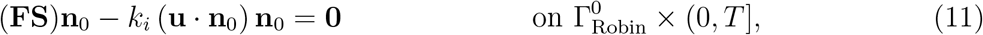

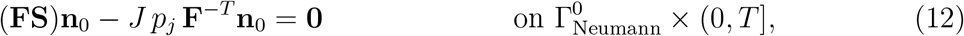

where **n**_0_ denotes the outward unit normal to *∂*Ω^0^, *k*_*i*_ is the spring stiffness associated with 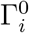, and *p*_*j*_ is the prescribed endocardial blood pressure acting on each 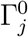.

Eq. 11 is a Robin boundary condition applied to the venous openings 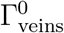, the valve annuli 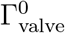, and the pericardial surfaces of the right atrium 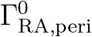 and left atrium 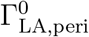. These regions are illustrated in Fig 6. The values *k*_*i*_ associated with each region are reported in Table 4.

**Table 4:** Spring stiffness coefficients used in the Robin boundary condition (11).

| Region | $\Gamma^0_{\text{veins}}$ | $\Gamma^0_{\text{valve}}$ | $\Gamma^0_{\text{RA,peri}}$ | $\Gamma^0_{\text{LA,peri}}$ |
| --- | --- | --- | --- | --- |
| $k_i$ (kPa mm<sup>-1</sup>) | 10.0 | 0.5 | 0.6 | 0.25 |

**Figure 6:**
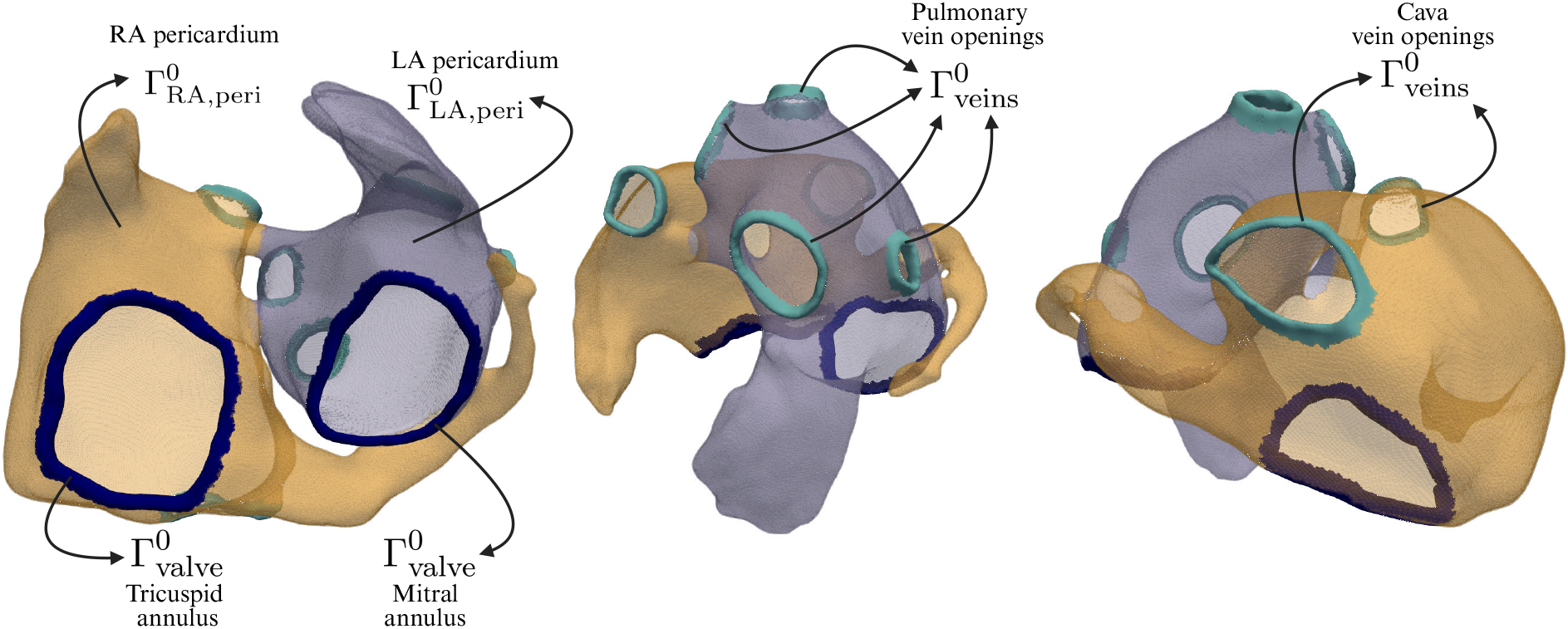
Spring boundary condition regions. Spring boundary conditions were applied to the atrioventricular valve annuli (mitral and tricuspid), venous openings (four pulmonary veins and the superior and inferior vena cava), left-atrial pericardium, and right-atrial pericardium.

The choice of these values was guided by previous modelling studies and physiological considerations:

- **Venous openings:** Imposing prescribed displacement (Dirichlet) boundary conditions at the atrial–venous junctions leads to unrealistic behaviour [11], as the atria tend to inflate against a rigid boundary. To allow more physiologically compliant deformation while preserving structural support at the venous junctions, a Robin (spring) boundary condition was adopted with a stiffness of *k* = 10.0 kPa mm^−1^.
- **Valve annuli:** The atrioventricular annuli (mitral and tricuspid) undergo physiological motion during the cardiac cycle, reflecting coupling between atrial and ventricular dynamics [80, 81, 82, 83]. To permit such motion, a moderate spring stiffness of *k* = 0.5 kPa mm^−1^ was assigned. This choice avoids artificially fixing the annuli and is consistent with experimental and clinical observations indicating that they are compliant, deformable structures rather than rigid boundaries [80, 83, 59, 84].
- **Pericardial surfaces:** The stiffness values associated with the right and left atrial pericardium were calibrated as part of the electromechanical calibration procedure to reproduce physiologically realistic atrial deformation.

Eq. 12 represents a pressure-type (Neumann) boundary condition that balances internal stresses with the blood pressure load applied at the endocardium of both atria (LA and RA). These endocardial pressures were provided through coupling with the 0D circulatory system model, ensuring physiologically consistent loading conditions throughout the cardiac cycle.

#### 2.2.3. Circulatory system

Circulatory system dynamics and cardiac hemodynamics were described using a 0D closed-loop model. The model reproduced hemodynamic pressures *p* and flows *Q* in the systemic and pulmonary circulations, as well as the chamber volumes *V* of the four cardiac cavities (LA, RA, left ventricle (LV), and right ventricle (RV)). The closed-loop circulation was formulated following established approaches proposed by Blanco *et al*. [66] and Regazzoni *et al*. [67], providing a compact yet physiologically consistent representation of the cardiac, systemic and pulmonary circuits suitable for coupling with the three-dimensional electromechanical atrial model.

Model parameters governing vascular properties were adjusted within physiological limits. The timing of the chamber elastance waveforms was adapted such that the onset of atrial contraction occurred at *t* = 0, in contrast to the ventricular-first timing assumed in [66], effectively delaying ventricular activation by 0.2 s. The elastance parameters were further tuned as part of the electromechanical calibration procedure to reproduce physiologically realistic hemodynamics.

A schematic of the circuit is shown in Fig 7. The full set of 0D model parameters is summarized in S3 Table.

**Figure 7:**
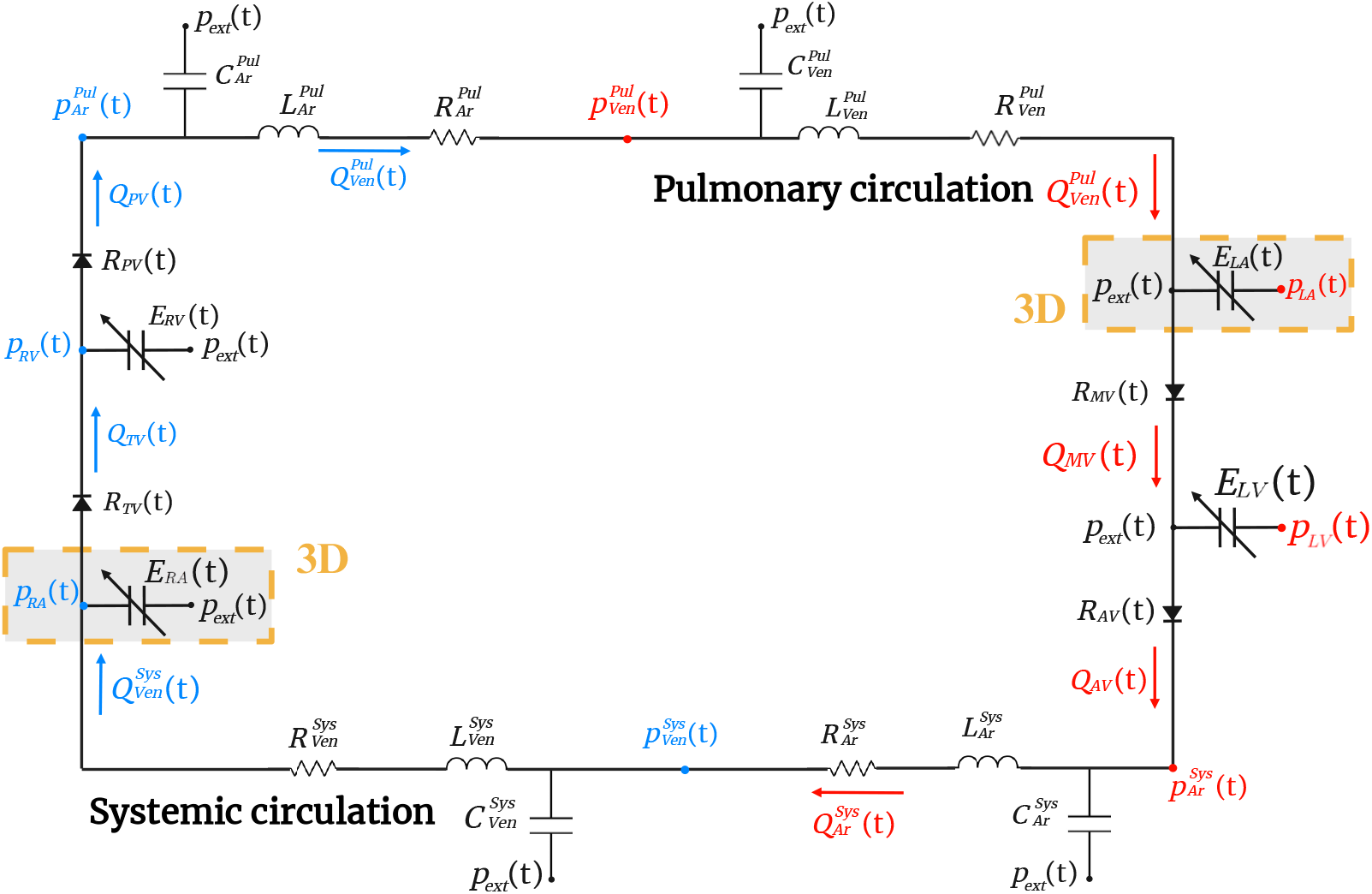
Closed-loop circulatory model diagram. The 0D closed-loop circuit is shown, where the 3D electromechanical atrial model replaces the components highlighted in the diagram.

### 2.3. Calibration of the model to reproduce physiological atrial function

#### Diffusion calibration

The longitudinal diffusion coefficient *D*_f_ is calibrated to ensure that simulated local activation times (LATs) reproduce physiological measurements. Under the anisotropic diffusion formulation introduced in Eq. 3, the calibration involves a single degree of freedom, with the transverse and normal diffusion components determined by the prescribed anisotropy ratios. Reference LATs were obtained from the work of Lemery *et al*. [85], who reported regional activation times in healthy subjects.

The calibration procedure consists of the following steps:

1. **Baseline simulation**. A six-beat simulation was performed with stimulation applied at the SAN using a basic cycle length (BCL) of 0.8 s, representative of resting sinus rhythm [86]. The first five beats were used to allow the system to reach a periodic steady state.
2. **Regional LAT extraction**. LATs were computed as the mean activation time within each anatomical region during the sixth simulated beat (*t* = 4.0–4.8 s).
3. **Diffusion tuning**. Regional values of *D*_f_ were manually adjusted until the simulated LATs fell within the reported physiological ranges. The regions used for calibration and the corresponding reference values are shown in Fig 8a.

**Figure 8:**
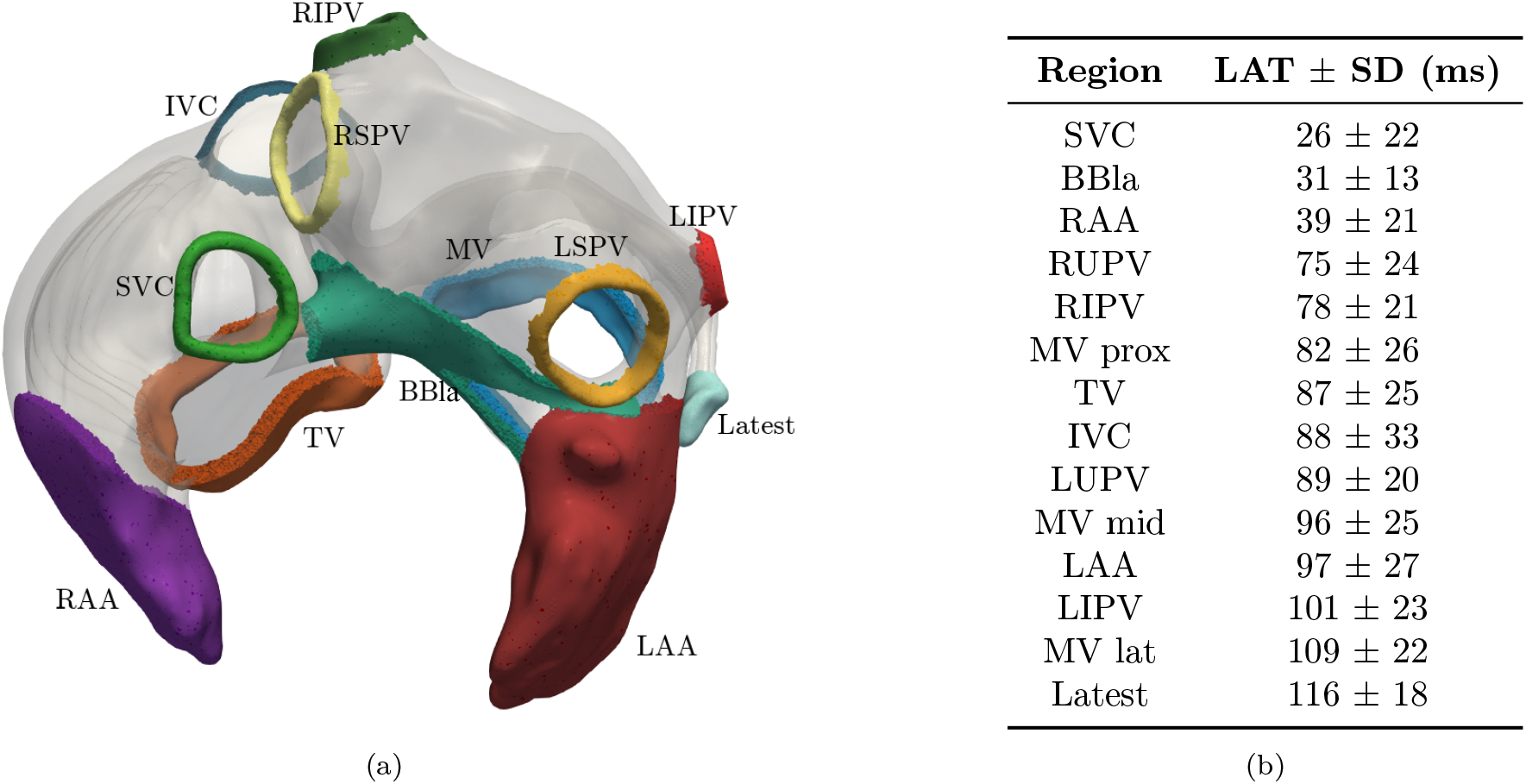
Atrial regions used to measure LAT for model calibration. (a) Anatomical locations where LATs were measured. (b) Reference LATs (mean ± SD) for the corresponding regions, reported by Lemery et al. [85].

#### 3D mechanical – 0D hemodynamic calibration

Following electrophysiological calibration, the electromechanical model was calibrated to reproduce physiological atrial function under sinus rhythm. As in the electrical diffusion calibration, simulations were run for six consecutive beats with a BCL of 0.8 s, and quantitative analysis was performed on the final beat after a periodic steady state was reached.

Calibration targeted clinically relevant macroscopic biomarkers. In particular, atrial minimum (*V*_min_), maximum (*V*_max_), and pre-atrial-contraction (*V*_preAC_) volumes, together with reservoir (EF_reser_), conduit (EF_cond_), and booster (EF_boost_) atrial EFs, were matched to reference values reported for a large cohort of healthy subjects (400 patients) by Gao *et al*. [87], considering only the male subcohort for consistency with the patient-specific anatomy. Phasic atrial EFs were computed following standard definitions [88]:

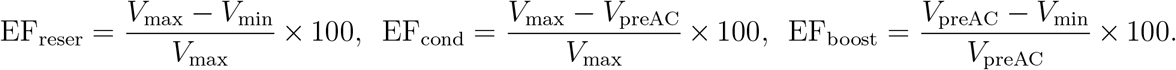

Only a limited subset of parameters was adjusted, selected based on preliminary simulation experiments showing a clear and consistent impact on atrial volumes and EFs, while all remaining parameters were kept fixed. The tuned parameters were: the reference active tension *T*_ref_ ; the pericardial spring stiffness coefficients *k*_RA,peri_ and *k*_LA,peri_; the ventricular active and passive elastances (*E*_LV,act_, *E*_RV,act_, *E*_LV,pass_, *E*_RV,pass_); the ventricular contraction and relaxation durations (*d*_LV,c_, *d*_RV,c_, *d*_LV,r_, *d*_RV,r_); and the systemic and pulmonary arterial resistances (*R*_SYS,AR_, *R*_PUL,AR_).

### 2.4. Sensitivity analysis

To assess the sensitivity of atrial mechanical function to key model parameters, a local sensitivity analysis was performed on the coupled 3D electromechanical–0D hemodynamic framework. Parameters were selected across different components of the model governing atrial contraction, passive mechanics, and ventricular loading. Specifically, the reference active tension (*T*_ref_), the isotropic passive stiffness (*a*), the fiber passive stiffness (*a*_*f*_), the pericardial stiffness (*k*_peri_), and the left and right active (*E*_LV,act_, *E*_RV,act_) and passive (*E*_LV,pass_, *E*_RV,pass_) ventricular elastances [66] were considered, resulting in a total of eight perturbed parameters.

For each parameter, three values were tested: the calibrated baseline value, a reduced value equal to half of the baseline, and an increased value equal to twice the baseline. All simulations were initialized from the same baseline configuration and run for four consecutive cardiac cycles to allow the model to reach a periodic regime. The fourth beat was used for analysis, as variations in atrial pressures and volumes between successive beats were below 2%.

Sensitivity was quantified using the same atrial volumetric and functional biomarkers employed in the electromechanical calibration, namely *V*_min_, *V*_max_, *V*_preAC_, and the reservoir, conduit, and booster atrial EFs. All biomarkers were computed for the left and right atria.

### 2.5. Application to a persistent AF scenario

The model is applied to a persistent AF scenario. For doing so the strategy consisted of the following steps:

1. **Electrical remodelling**. Atrial electrical remodelling associated with persistent AF was introduced by scaling selected ionic conductances at the cellular level, in line with previous computational studies [13, 27]. Specifically, *I*_K1_ was increased by a factor of 2.0 in both atria [28, 89]. *I*_to_ and *I*_Kur_ were reduced in an atrium-specific manner, to 0.55 and 0.40 of baseline values in the RA, and to 0.25 and 0.55 in the LA, respectively [23]. *I*_Ks_ was increased by a factor of 2.5 in the RA and 2.0 in the LA [23]. Finally, *I*_CaL_ was reduced to 0.35 of its baseline value in both atria [90, 91]. Within each atrium, the same remodelling pattern was applied uniformly across all corresponding anatomical regions.
2. **Stabilization**. A five-beat stabilization in sinus rhythm was performed with a single stimulus applied at the SAN and a BCL of 0.8 s. Electrical remodelling was already active during this phase so that the 3D model reached a physiological steady state.
3. **Selection of ectopic site**. A single ectopic focus was placed near atrial regions known to exhibit high vulnerability to fibrillation, in this case adjacent to the PVs (Fig 9a) [92, 93].
4. **Determination of ectopic timing**. The timing of the ectopic stimulus was determined based on local recovery of excitability at the selected site, following a protocol analogous to a classical S1–S2 pacing protocol consisting of regular pacing beats (S1) followed by a single premature stimulus (S2). Recovery of excitability was estimated from the restitution behaviour of the fast sodium current inactivation gates *h* and *j* [94]. Excitability was assumed to be restored when the product *h · j* exceeded 0.1, as illustrated in Fig 9b.
5. **Ectopic stimulation**. A spherical ectopic stimulus of diameter 8 mm was applied at the moment the tissue regained excitability, as identified from the restitution curves of the sodium channel gates.
6. **AF induction and free evolution**. Following ectopic stimulation, SAN pacing was interrupted and the simulation was allowed to evolve freely. This reflects the clinical observation that, once AF is initiated, sinus rhythm is overridden by the arrhythmia and atrial activation becomes dominated by self-sustained fibrillatory activity [20]. Ventricular activation in the 0D elastance-based model continued to be prescribed with the same sinus rhythm as before and was used to assess AF persistence [20, 7, 69].

**Figure 9:**
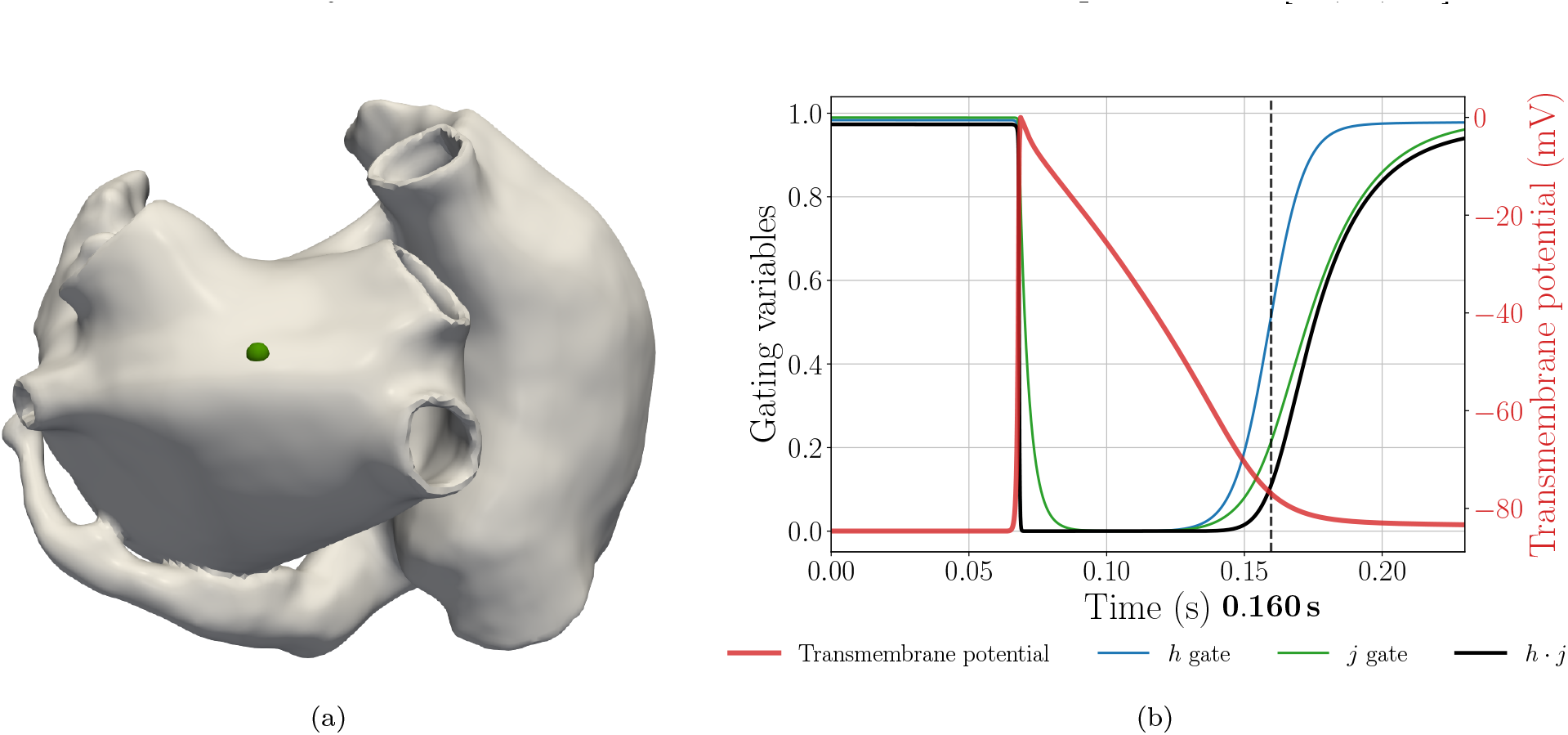
Workflow for AF induction through an ectopic trigger. (a) Location of the ectopic focus near the PVs. (b) Restitution behaviour of the sodium channel inactivation gates *h* and *j*, their product *h · j*, and the local transmembrane potential at the ectopic site. Recovery of excitability is identified from the evolution of *h · j*, with the dashed line indicating the time at which the excitability threshold is exceeded.

### 2.6. Computational aspects

Simulations were performed in an HPC environment due to the computational cost of resolving a fully coupled 3D electromechanical–0D hemodynamic heartbeat. All simulations were conducted using Alya [95, 56, 96, 68, 97], a finite element method (FEM) multiphysics simulation engine developed at the Barcelona Supercomputing Center (BSC) and ELEM Biotech S.L., specifically designed for large–scale strongly coupled problems.

The monodomain equation (Eq. 1) was discretized in space using linear finite elements. Time integration of the reaction–diffusion system was performed using an implicit Euler scheme for the diffusion term, while the ionic model was integrated using a Rush–Larsen method for the gating variables, exploiting their exponential formulation to improve numerical stability in the presence of stiff transmembrane dynamics and fibrillatory activation patterns. The resulting linear systems were solved using a conjugate gradient method with diagonal preconditioning. A time step of 5 × 10^−6^ s was used to accurately resolve the fast transmembrane dynamics.

The nonlinear momentum balance equation (Eq. 4) governing tissue deformation was discretized using linear finite elements. Time integration was performed using an implicit damped Newmark-beta scheme. The resulting nonlinear system, arising from the constitutive response and active stress formulation, was solved using a Newton–Raphson iterative procedure. The linearized systems were solved using GMRES with block, diagonal, and RAS preconditioning. A time step of 2 × 10^−5^ s was employed, consistent with the smoother temporal evolution of tissue deformation.

Electromechanical coupling was handled through a multi-instance strategy in which electrophysiology and solid mechanics were solved on separate meshes using distinct Alya instances, each responsible for its own geometric and physical domain. The two instances were coupled through a staggered communication scheme that allowed each physics to advance with its own time step. Coupling fields, namely intracellular calcium concentration and tissue displacement, were exchanged every four electrophysiology time steps, ensuring synchronization between electrical activation and mechanical contraction while preserving computational efficiency.

The interaction between both solvers was managed through the Dynamic Load Balancing (DLB) library [98, 99], which optimized CPU utilization by oversubscribing processes from the two Alya instances and improving intra-node communication through shared memory. This strategy enabled efficient concurrent execution of the electromechanical problem.

The right and left atria were modelled in 3D, whereas the ventricles and systemic and pulmonary circulations were represented using the 0D closed-loop formulation. Coupling between atrial mechanics and the circulation model was enforced through the Neumann boundary condition applied at the atrial endocardial surfaces (Eq. 12). Atrial volumes acted as Lagrange multipliers, ensuring consistency between 3D chamber deformation and the 0D hemodynamic state. The augmented system resulting from the Lagrange multipliers was handled using a Schur complement approach [100, 67] and adapted here to the biatrial configuration.

All simulations were executed on MareNostrum V^6^ at BSC. Each simulation used 560 CPU cores and required approximately 3 hours and 6 minutes to simulate one full electromechanical heartbeat.

## 3. Results

### 3.1. Calibration of the model to reproduce physiological atrial function

#### 3.1.1. Diffusion calibration

Region-specific diffusion parameters were calibrated to reproduce physiological atrial activation patterns. The resulting region-specific diffusion values are reported in Table 5.

**Table 5:** Electrical diffusion parameters obtained after calibration.

| Region | RA | CT | PVs | PMs | ISTH | SAN | FO | CS | LA | FO limb | BBla |
| --- | --- | --- | --- | --- | --- | --- | --- | --- | --- | --- | --- |
| $D_f$ (cm<sup>2</sup>/s) | 1.80 | 8.00 | 1.10 | 10.00 | 1.00 | 0.10 | 0.01 | 15.00 | 1.95 | 5.00 | 4.00 |
| $D_t, D_n$ (cm<sup>2</sup>/s) | 0.63 | 1.20 | 0.55 | 1.50 | 1.00 | 0.10 | 0.01 | 7.50 | 0.6825 | 0.75 | 0.60 |

The impact of these calibrated diffusion values on atrial electrical activation is illustrated in Fig 10, which reports simulated LATs across the atrial regions listed in Fig 8. After calibration, simulated regional LATs fall within the physiological ranges reported in the literature [85]. In contrast, a simulation setting a homogeneous diffusion tensor, **D** = diag(1.8, 0.63, 0.63) cm^2^*/*s, exhibit pronounced regional activation delays, with deviations exceeding 40 to 70 ms in several atrial regions.

**Figure 10:**
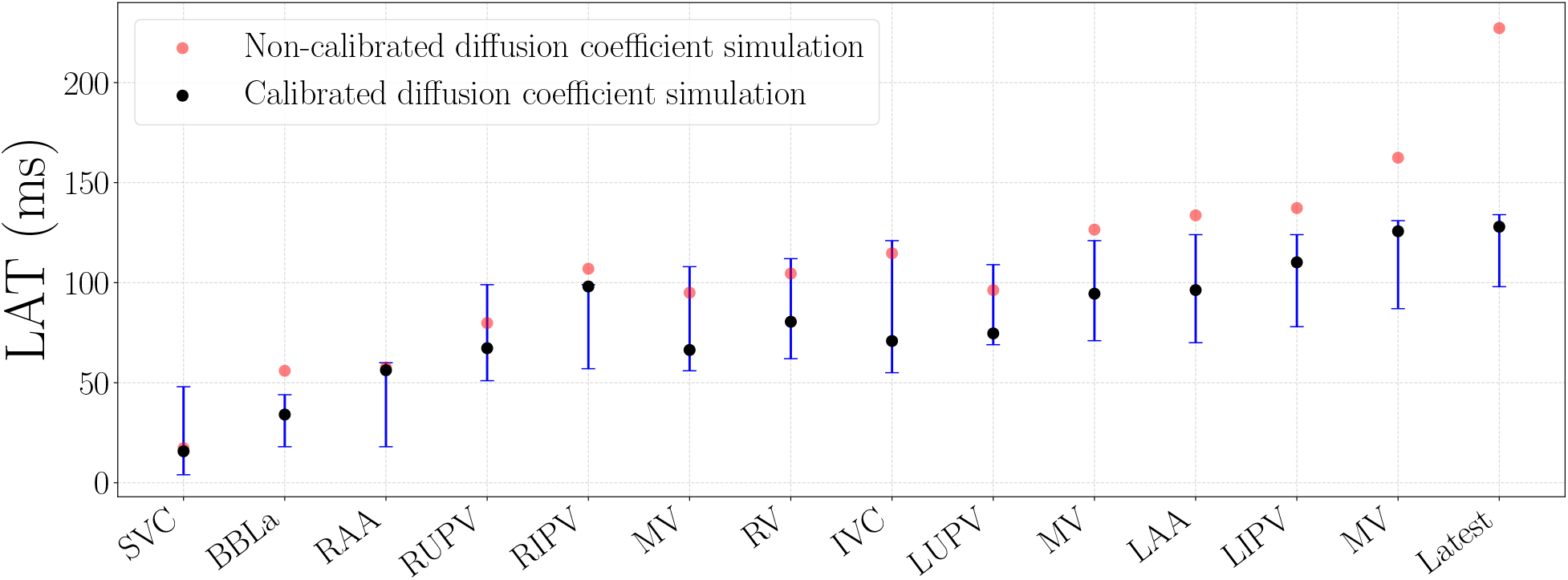
Diffusion calibration of regional LATs. Simulated LATs during the sixth heartbeat (*t* = 4.0–4.8 s) are shown for all atrial regions reported in Fig 8 using calibrated diffusion parameters (black markers). For comparison, results obtained with a homogeneous diffusion tensor are shown in red, and reference physiological LAT ranges from Lemery *et al*. [85] are shown in blue.

#### 3.1.2. 3D mechanical–0D hemodynamic calibration

Fig 11 summarizes atrial mechanical biomarkers obtained after the electromechanical calibration. The calibrated simulation reproduces all reference atrial volumes and phasic EFs within the physiological ranges reported by Gao *et al*. [87].

**Figure 11:**
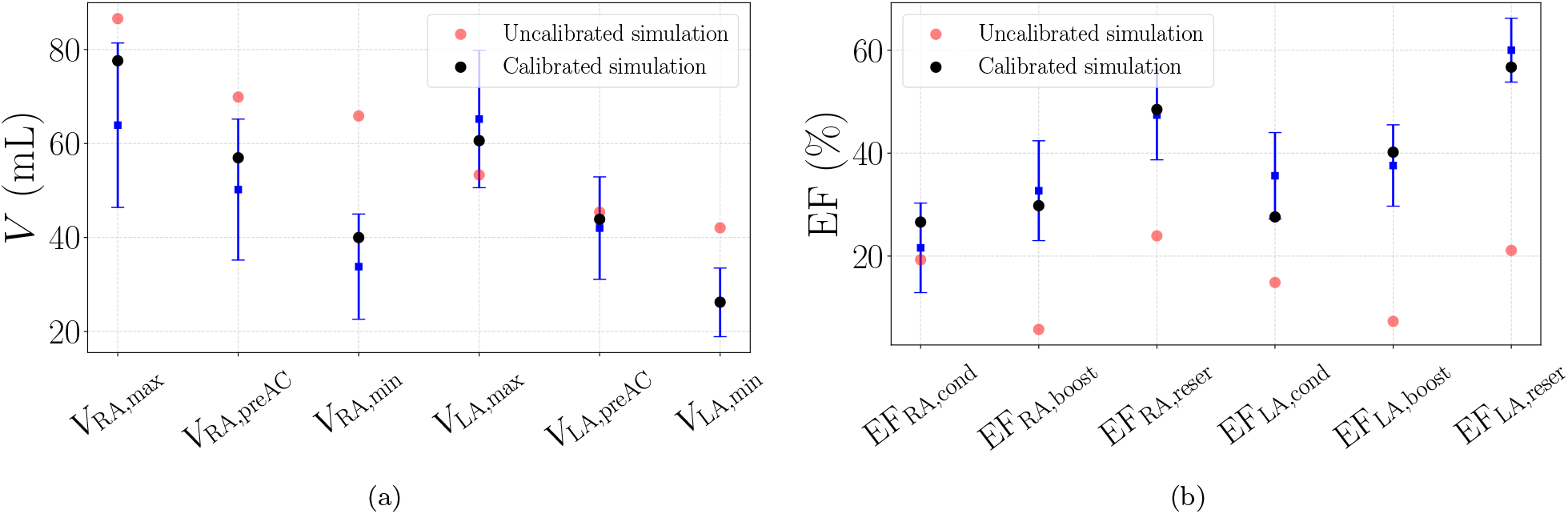
Calibration of atrial mechanical biomarkers. Comparison between simulated atrial mechanical biomarkers and physiological reference values from Gao *et al*. [87]. Calibrated electromechanical results are shown in black, uncalibrated results in red, and reference values (mean ± SD) in blue. (a) Atrial volumes (*V*_min_, *V*_max_, *V*_preAC_). (b) Reservoir, conduit, and booster atrial EFs.

For comparison, results from an uncalibrated electromechanical simulation (red markers), in which parameters were left at literature reference values, are also shown. The calibrated and uncalibrated parameter values are listed in Table 6. The uncalibrated case exhibits marked deviations from physiological ranges, reaching differences of up to 30 mL in atrial volumes and approximately 50% in atrial EFs. This contrast highlights the necessity of consistent 3D mechanical and 0D hemodynamic parameter tuning to achieve physiologically realistic atrial function.

**Table 6:** Parameters used in the 3D electromechanical–0D hemodynamic calibration.

| Parameter | Symbol | Unit | Reference value | Calibrated value |
| --- | --- | --- | --- | --- |
| Bulk modulus | $K$ | kPa | – | 1218.5 |
| Reference tension | $T_{\text{ref}}$ | kPa | 120 | 720 |
| LV active elastance | $E_{\text{LV,act}}$ | mmHg/mL | 2.75 | 3.75 |
| LV passive elastance | $E_{\text{LV,pass}}$ | mmHg/mL | 0.08 | 0.07 |
| RV active elastance | $E_{\text{RV,act}}$ | mmHg/mL | 0.55 | 0.60 |
| RV passive elastance | $E_{\text{RV,pass}}$ | mmHg/mL | 0.05 | 0.04 |
| LV contraction onset | $t_{\text{LV,c}}$ | s | 0.00 | 0.20 |
| RV contraction onset | $t_{\text{RV,c}}$ | s | 0.00 | 0.20 |
| LV contraction duration | $d_{\text{LV,c}}$ | s | 0.34 | 0.45 |
| RV contraction duration | $d_{\text{RV,c}}$ | s | 0.34 | 0.41 |
| LV relaxation duration | $d_{\text{LV,r}}$ | s | 0.15 | 0.22 |
| RV relaxation duration | $d_{\text{RV,r}}$ | s | 0.15 | 0.22 |
| Systemic arterial resistance | $R_{\text{SYS,AR}}$ | mmHg·s/mL | 0.8000 | 0.4500 |
| Pulmonary arterial resistance | $R_{\text{PUL,AR}}$ | mmHg·s/mL | 0.1625 | 0.0280 |

### 3.2. Electromechanical atrial dynamics after calibration

#### 3.2.1. Spatiotemporal electromechanical dynamics

Fig 12 illustrates some representative snapshots of the sixth simulated heartbeat ([4.0– 4.8] s) of the fully calibrated case, showing the propagation of the transmembrane potential within the deforming biatrial geometry. Between 4.01 and 4.24 s, the atria underwent active contraction, reaching their minimum volumes (booster phase). This was followed by atrial filling during ventricular systole between 4.24 and 4.57 s (reservoir phase). Subsequently, passive emptying into the ventricles occurred during early ventricular diastole between 4.57 and 4.8 s (conduit phase).

**Figure 12:**
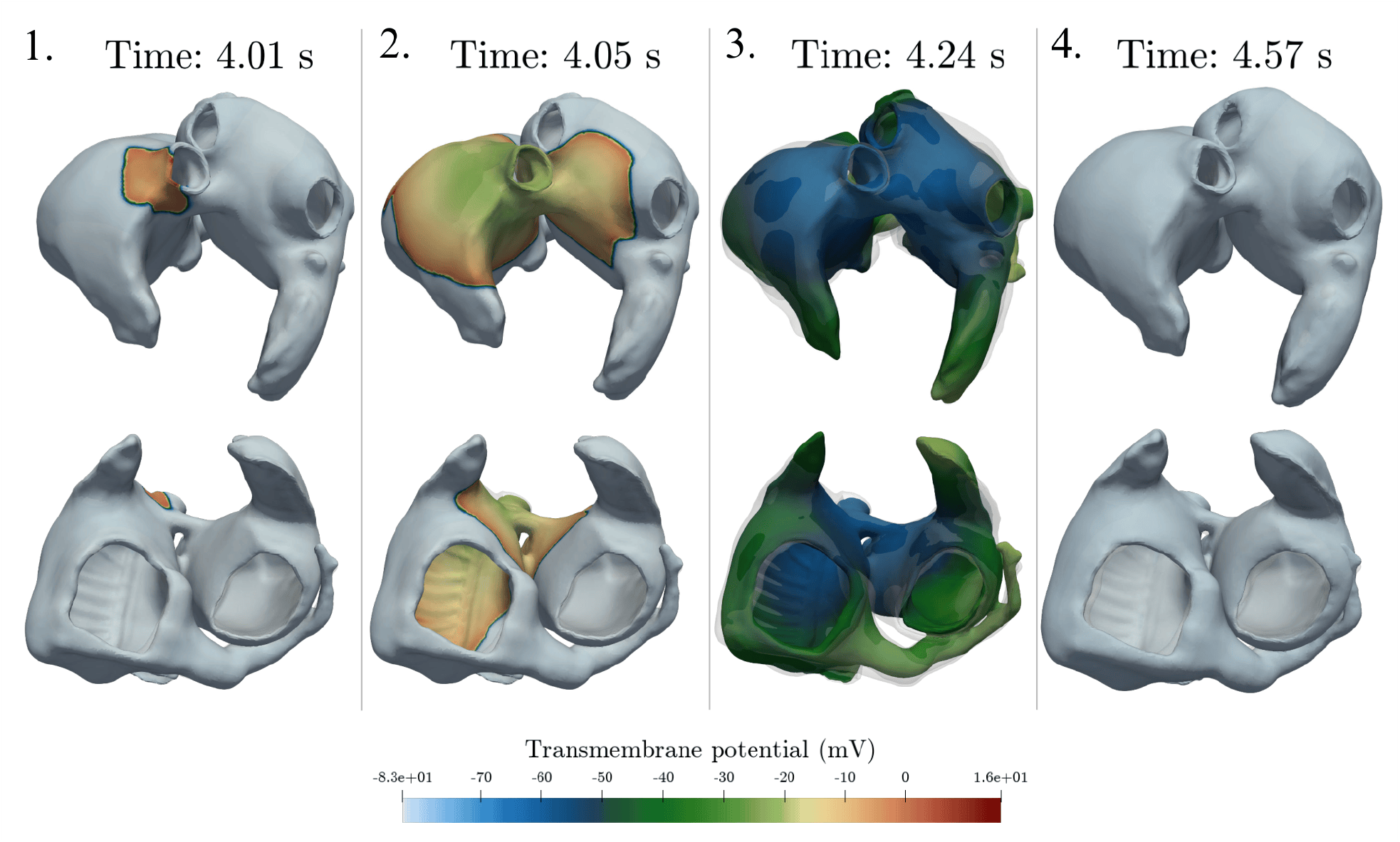
Electromechanical activation snapshots. Transmembrane potential propagation through the 3D biatrial anatomy is shown together with tissue deformation relative to the initial configuration (semitransparent surface).

#### 3.2.2. Atrial pressure–volume loops

Fig 13 presents the pressure–volume (PV) loops of the left and right atria together with their corresponding pressure-and volume-time traces for the sixth beat of the fully calibrated simulation. Both atria display a similar *figure-eight* pattern, in which two loops can be clearly identified: the A-loop (Atrial-loop), associated with active atrial contraction (booster pump) and the early reservoir phase and the V-loop (Ventricular filling loop), associated with passive reservoir and conduit behaviour.

**Figure 13:**
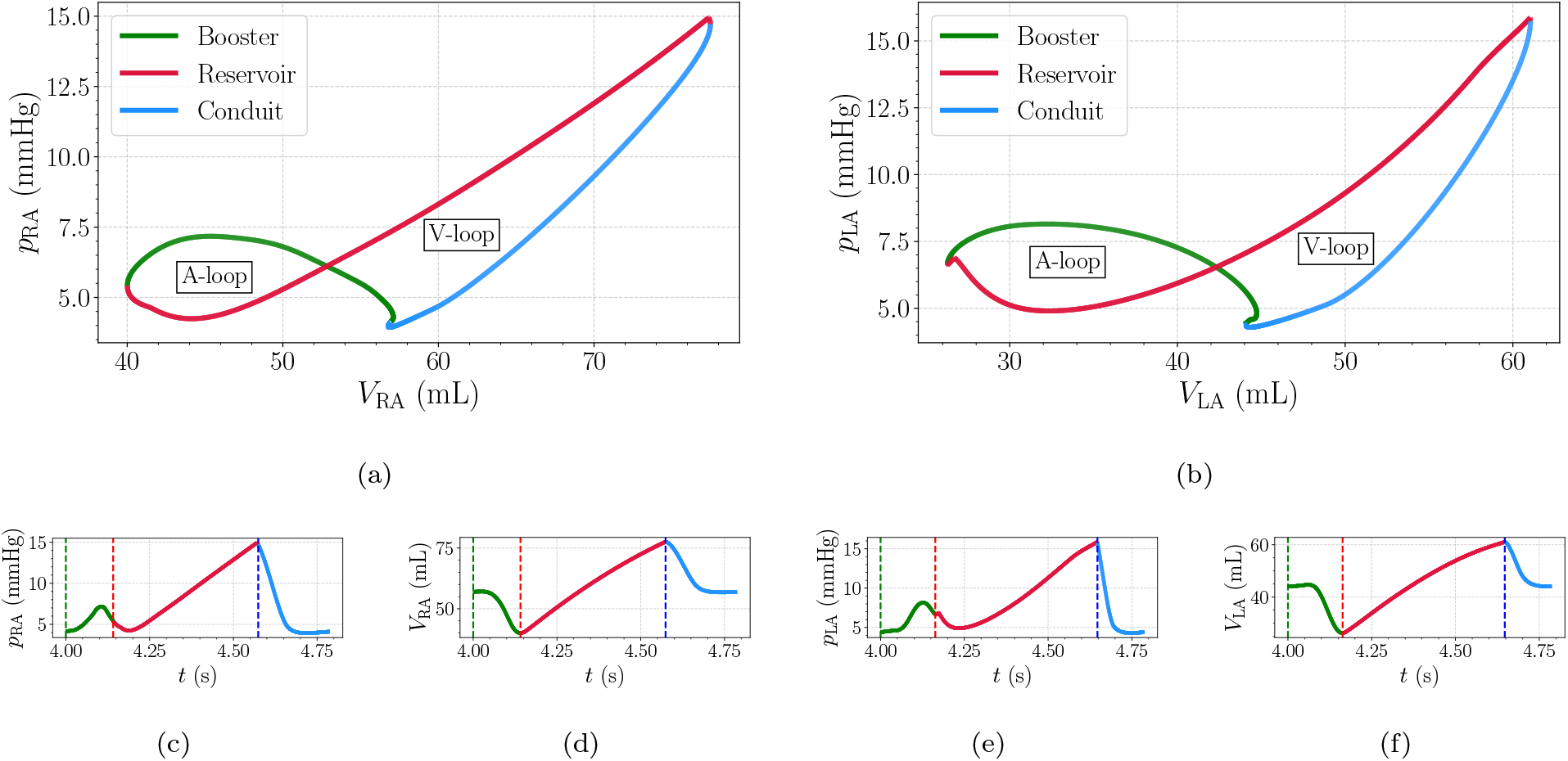
Atrial pressure–volume loops and temporal traces. Results correspond to the sixth simulated beat of the fully calibrated case. The reservoir, conduit, and booster phases are identified across pressure–volume loops and temporal traces. (a) Right atrial pressure–volume loop with V-loop and A-loop annotations. (b) Left atrial pressure–volume loop. (c) Right atrial pressure as a function of time with vertical lines marking the transitions between atrial phases. (d) Right atrial volume as a function of time. (e) Left atrial pressure trace. (f) Left atrial volume trace.

For comparison, Fig 14 also shows the atrial pressure–volume loops obtained from the uncalibrated electromechanical simulation. In this case, the pressure–volume loops do not display a figure-eight pattern. The initial portion of the loop is characterized by a pronounced increase in pressure with limited variation in volume, followed by subsequent pressure and volume changes that do not form two clearly separated loops. As a result, the PV trajectories do not exhibit a distinct A-loop and V-loop structure over the cardiac cycle.

**Figure 14:**
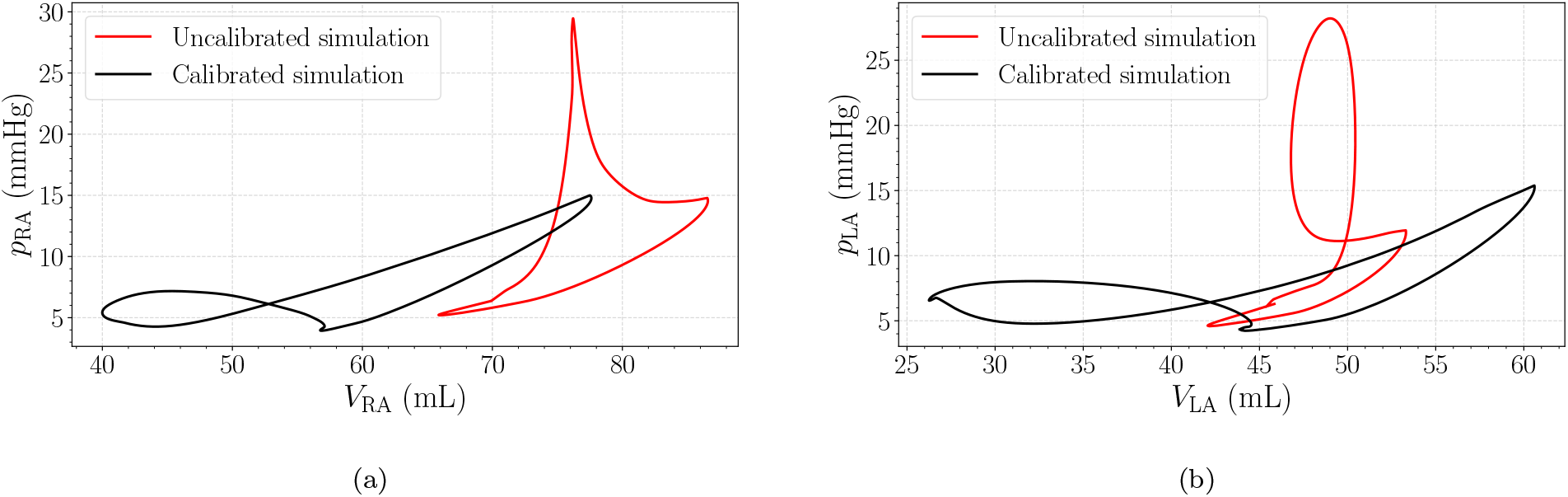
Comparison of atrial pressure–volume loops. Pressure–volume loops of the sixth simulated beat are shown for the calibrated (black) and non-calibrated (red) cases for (a) the right atrium and (b) the left atrium.

### 3.3. Sensitivity analysis

The sensitivity analysis reveals parameter-specific effects on atrial and ventricular pressure–volume behaviour and phasic mechanical biomarkers.

#### 3.3.1. Reference active tension T_ref_

As shown in Fig 15, increasing the reference active tension produced a leftward shift in both minimum and maximum atrial volumes, while pressures remained within a similar range. The pre-atrial-contraction volume remained nearly unchanged across the tested values. The functional biomarkers reflected these changes: the booster EF increased clearly with *T*_ref_ (4%), while the reservoir and conduit components varied only slightly (1% and 0.5% respectively). The overall atrial loop morphology and timing remained qualitatively similar, while ventricular pressure–volume loops did not show significant changes across the tested values.

**Figure 15:**
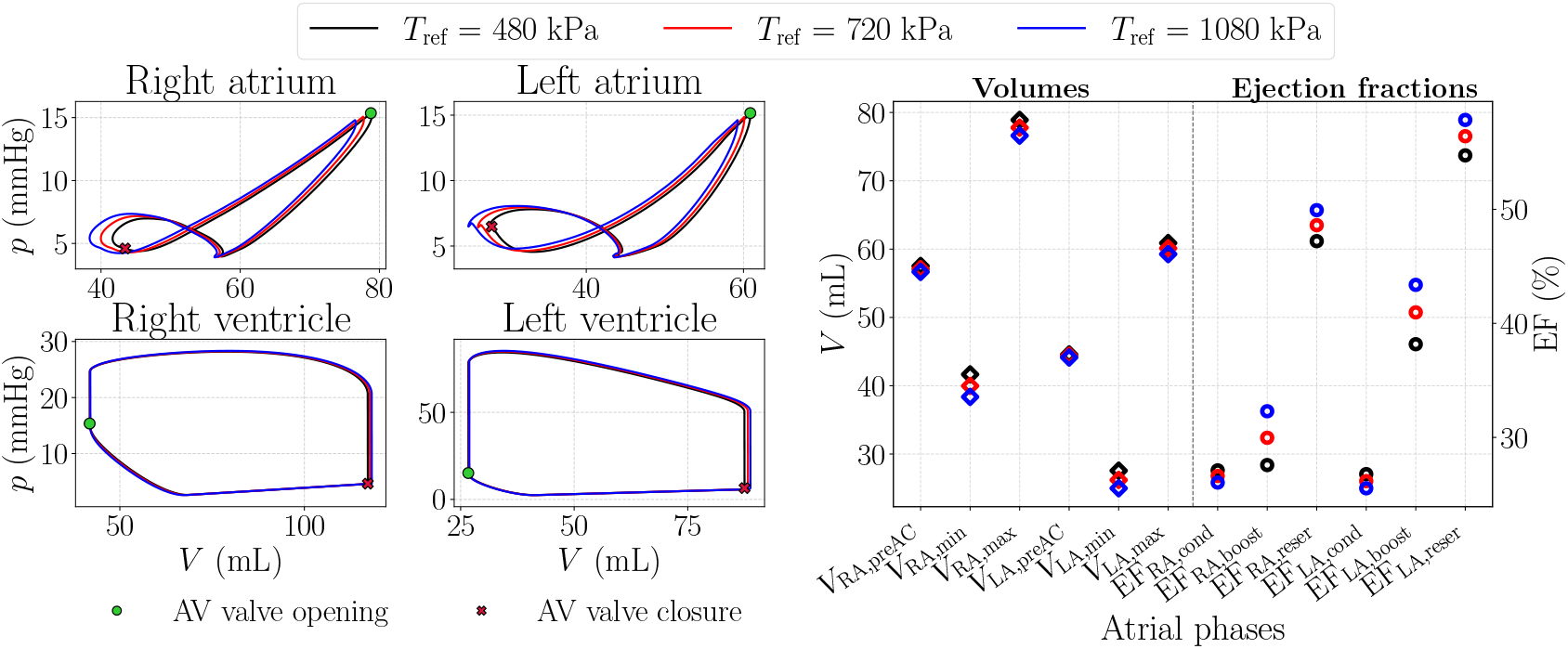
Effect of varying the reference active tension *T*_ref_. Atrial pressure–volume loops for the tested parameter values, with atrioventricular valve opening and closure shown for comparison with ventricular loops, together with the corresponding atrial volumes and EFs.

#### 3.3.2. Isotropic ground-matrix stiffness a and fiber stiffness a_f_

As shown in Fig 16, scaling the isotropic ground-matrix stiffness *a* and the fiber stiffness *a*_f_ produced consistent shifts in the atrial pressure–volume loops. In both cases, increasing stiffness reduced the overall volume range, primarily through a decrease of 2 mL in the maximum and pre-atrial-contraction volumes, while the minimum volume exhibited smaller changes (*<* 0.01%). The associated functional biomarkers followed the same trends, with noticeable variations in reservoir-related quantities and comparatively modest variations in the booster component. The overall atrial loop morphology remained qualitatively similar, and ventricular pressure–volume loops did not exhibit significant changes within the explored stiffness range.

**Figure 16:**
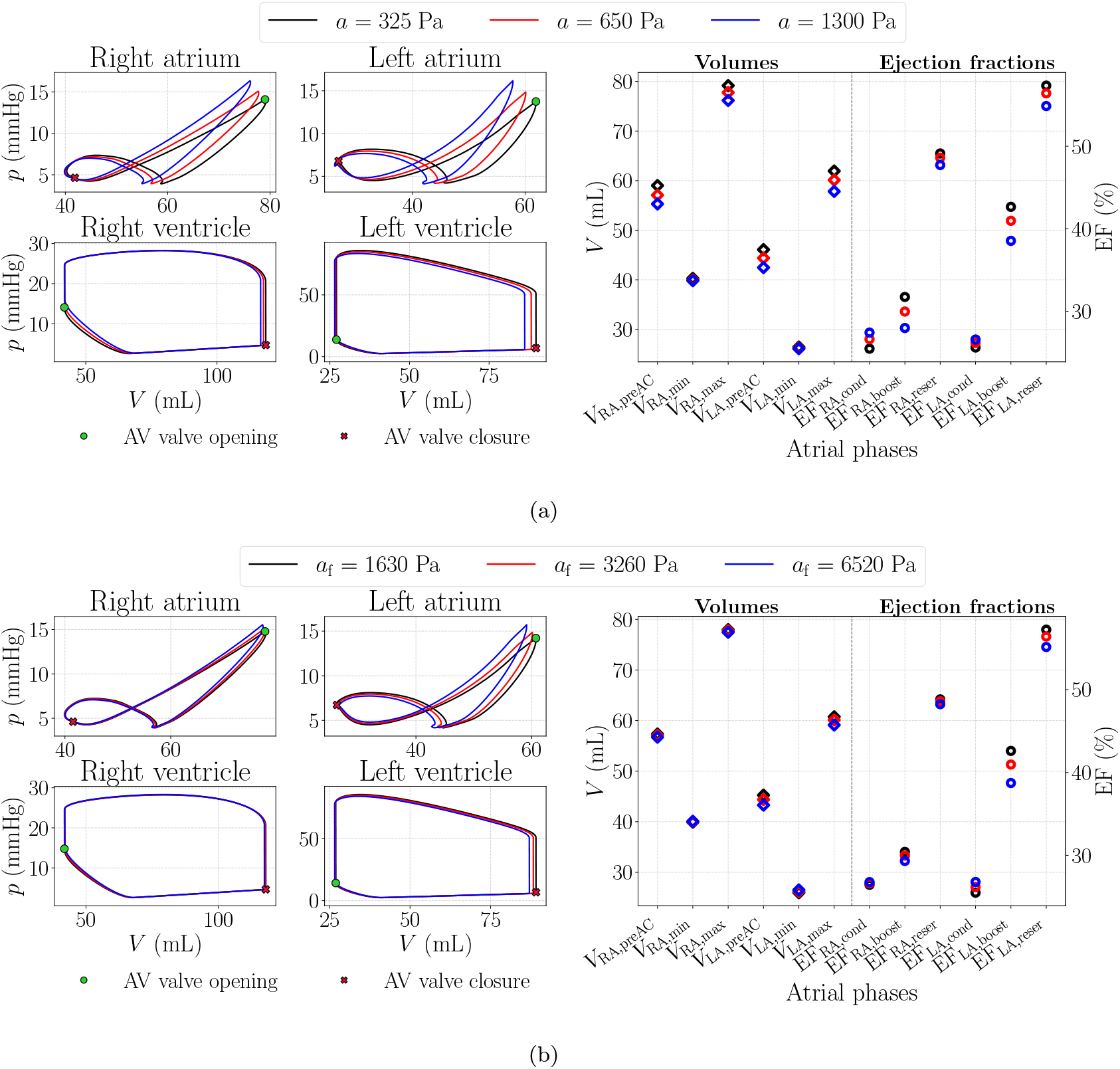
Effect of varying passive stiffness parameters. (a) Influence of the isotropic ground-matrix stiffness *a*. (b) Influence of the fiber stiffness *a*_f_. Each panel shows atrial pressure–volume loops for the tested parameter values, with atrioventricular valve opening and closure indicated for comparison with ventricular loops (left), together with the corresponding atrial volumes and EFs (right).

The effects of varying *a*_f_ were qualitatively similar to those observed for *a*, but less pronounced in magnitude (maximum and pre-atrial-contraction variations only reach 1 mL in magnitude).

#### 3.3.3. Spring stiffness coefficient of the atrial pericardium k_peri_

As shown in Fig 17, increasing the pericardial spring stiffness coefficient reduced both the maximum and pre-atrial-contraction volumes (5 mL and 2 mL respectively), while the minimum volume remained nearly unchanged. The ventricular loops also showed smaller volume excursions for higher stiffness values. The functional biomarkers followed these trends: the reservoir and booster EFs decreased with increasing stiffness, whereas the conduit fraction varied only slightly. The overall atrial loop morphology remained qualitatively similar across the tested range, and ventricular pressure–volume loops did not show significant changes.

**Figure 17:**
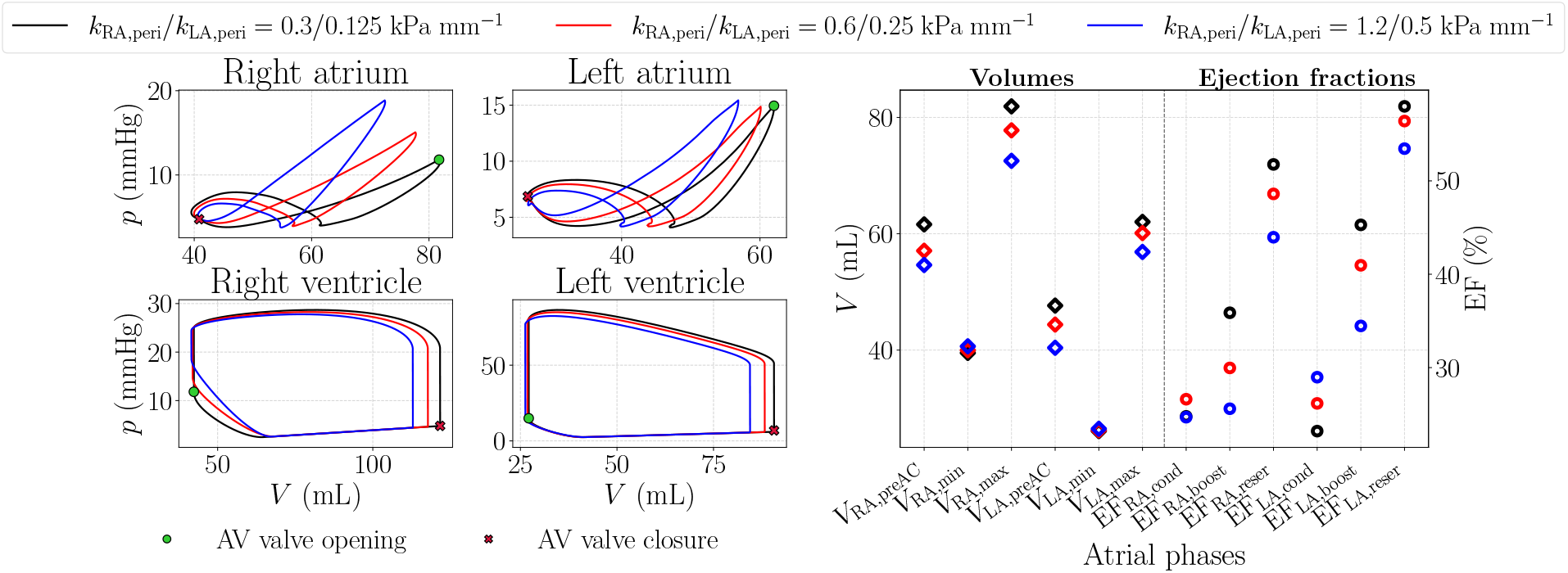
Effect of varying pericardial spring stiffness *k*_peri_. Atrial pressure–volume loops for the tested parameter values, with atrioventricular valve opening and closure shown for comparison with ventricular loops, together with the corresponding atrial volumes and EFs.

#### 3.3.4. Ventricular active elastances E_act_

As shown in Fig 18, increasing the left-ventricular active elastance *E*_LV,act_ primarily affected the left-sided chambers. The left-atrial pressure–volume loop shifted downward during the booster phase, accompanied by a reduction in the pre-atrial-contraction volume of approximately 2 mL, while the right-atrial loop remained largely unchanged. These changes reduced the enclosed area of the atrial A-loop, which is interpreted as a measure of the work done by the atria during active contraction [101]. At the ventricular level, higher values of *E*_LV,act_ resulted in a reduced left-ventricular volume excursion. Consequently, the left-ventricular EF increased from 56.02% (black) to 69.76% (red) and 79.08% (blue).

**Figure 18:**
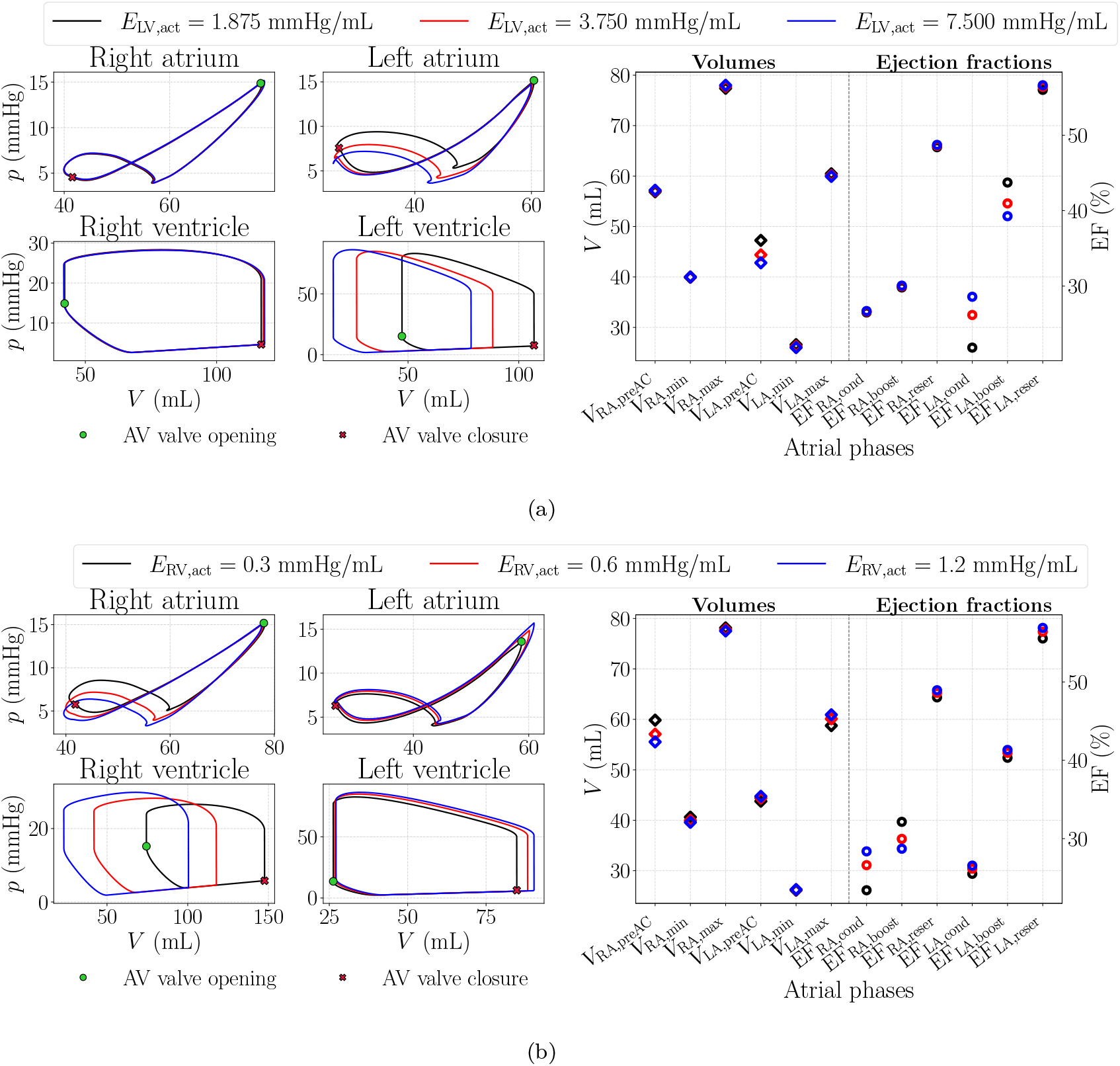
Effect of varying the active ventricular elastance parameters. (a) Influence of the leftventricular active elastance *E*_LV,act_. (b) Influence of the right-ventricular active elastance *E*_RV,act_. Each panel shows atrial pressure–volume loops for the tested parameter values, with atrioventricular valve opening and closure indicated for comparison with ventricular loops (left), together with the corresponding atrial volumes and EFs (right).

Analogously, variations in the right-ventricular active elastance *E*_RV,act_ mainly influenced the right-sided chambers, inducing comparable changes in the right atrium and right ventricle, while exerting only minor effects on the left-sided dynamics. In both ventricles, the changes in active elastance were consistently reflected in atrial functional biomarkers: the reservoir EF remained nearly constant (variation *<* 0.01%), the conduit fraction showed a modest increase (3%), and the booster fraction decreased with increasing ventricular active elastance (2%).

#### 3.3.5. Ventricular passive elastances E_pass_

As shown in Fig 19, increasing the left-ventricular passive elastance *E*_LV,pass_ predominantly affected the left-sided chambers. Left-atrial pressures increased during the booster and early reservoir phases (7 mmHg), and the left-atrial pressure–volume loop shifted toward higher pressure values during the booster phase, together with a marked increase in the pre-atrial-contraction volume (6 mL). In contrast to the active elastance case, these changes resulted in an enlargement of the atrial A-loop area, indicating an increase in the work performed by the left atrium during atrial contraction. The right atrium exhibited only minor variations (*<* 0.01 %). At the ventricular level, higher *E*_LV,pass_ led to an upward shift of the left-ventricular pressure–volume loop, accompanied by a reduction in the ventricular volume range.

**Figure 19:**
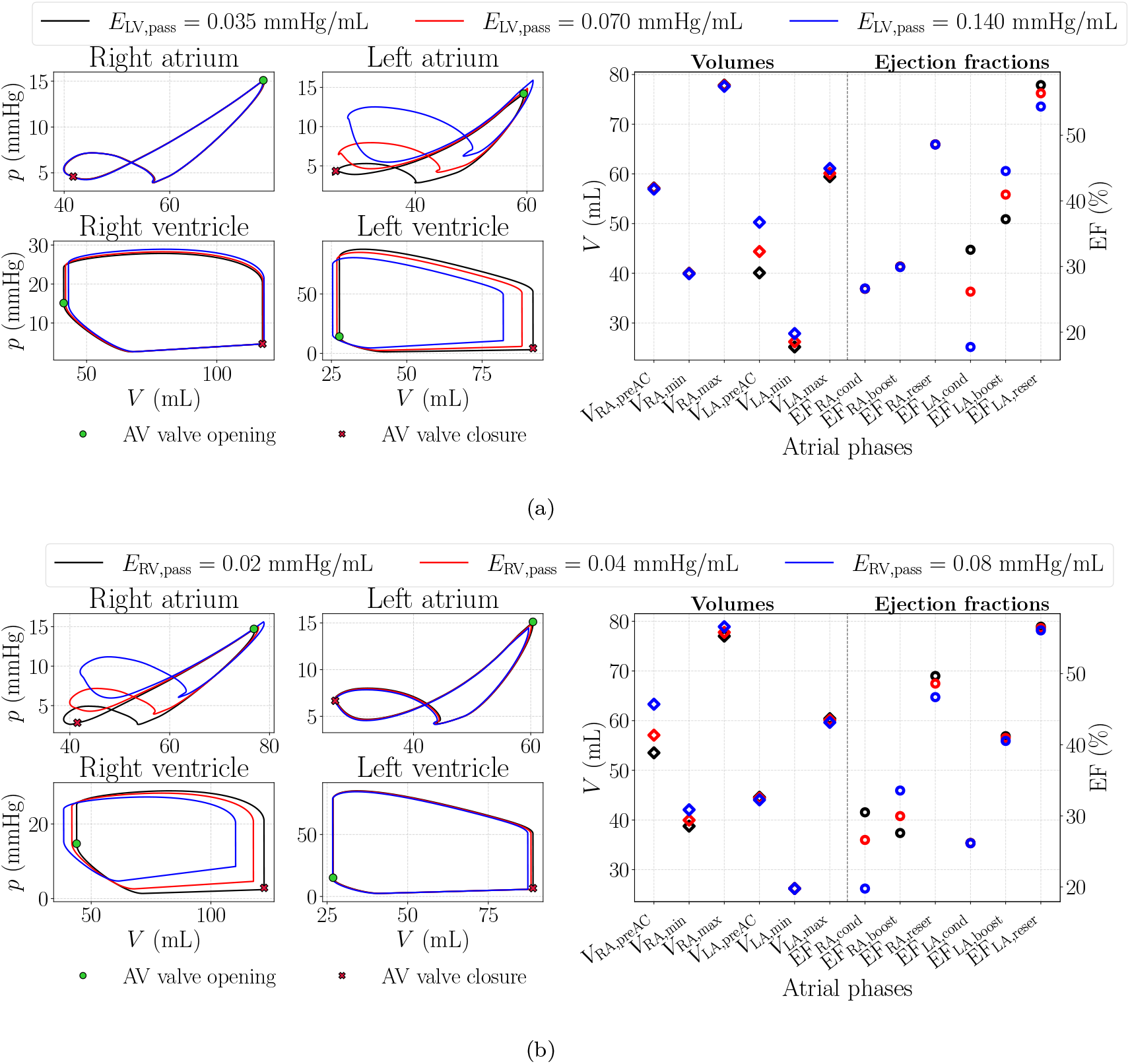
Effect of varying the passive ventricular elastance parameters. (a) Influence of the left-ventricular passive elastance *E*_LV,pass_. (b) Influence of the right-ventricular passive elastance *E*_RV,pass_. Each panel shows atrial pressure–volume loops for the tested parameter values, with atrioventricular valve opening and closure indicated for comparison with ventricular loops (left), together with the corresponding atrial volumes and EFs (right).

Varying the right-ventricular passive elastance *E*_RV,pass_ produced an analogous pattern on the right side, with corresponding changes in the right atrium and right ventricle and limited influence on the left chambers. In both cases, the functional biomarkers followed similar trends: the reservoir EF decreased with passive elastance (2%), the conduit fraction largely decreased (10%), and the booster fraction rised (3%).

### 3.4. Application to a persistent AF scenario

#### 3.4.1. AF initiation

The S1–S2 protocol successfully induced a sustained episode of AF. In this simulation, a single premature ectopic stimulus was delivered at *t* = 4.16 s, corresponding to the time at which local tissue excitability had recovered at the selected ectopic site. This perturbation initiated a sequence of events that led to reentrant activity in the LA. The sequence of events leading to reentry is illustrated in Fig 20.

**Figure 20:**
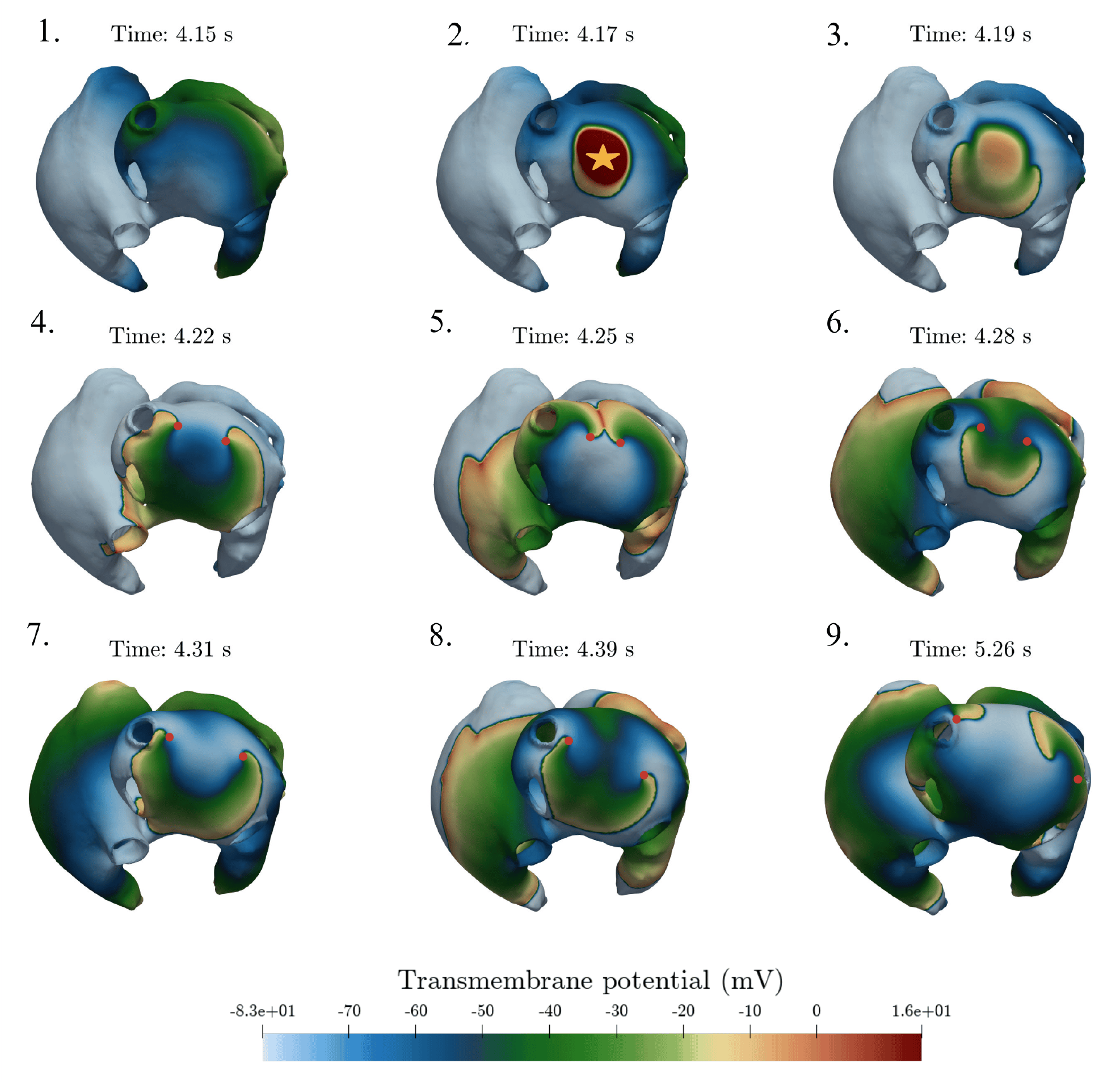
Initiation of atrial fibrillation by ectopic stimulation. A premature ectopic stimulus applied at *t* = 4.16 s triggers unidirectional conduction block and subsequent reentry in the left atrium, leading to the formation of two counter-rotating rotors. The ectopic stimulation site is indicated in the figure, and rotor cores are marked in the final snapshots.

The final stabilization beat initiated at *t* = 4.0 s was completing its propagation immediately before stimulus delivery (*t* = 4.15 s). In particular, regions of the LA close to the PVs had not fully recovered, resulting in spatial differences in tissue excitability. When the ectopic stimulus was delivered, it activated a small localized area but led to asymmetric propagation: the resulting wavefront travelled toward excitable tissue in the anterior LA, while propagation in the opposite direction was prevented by tissue that remained refractory. This unidirectional conduction block was clearly visible by *t* = 4.19 s.

After this initial block, the wavefront propagated through the surrounding excitable tissue and gradually bent around the region activated by the ectopic stimulus, which remained partially inexcitable. As the wave moved forward and the tissue behind it recovered, a gap in conduction appeared, allowing the curved wavefront to turn back and re-enter the area it had just crossed. Between *t* = 4.22 s and *t* = 4.25 s, this mechanism led to the initiation of reentry, observed as the formation of two spiral waves in the LA.

The locations around which these spiral waves rotated correspond to the rotor cores shown in Fig 20. By *t* = 4.28 s, the rotors were fully established and exhibited stable rotation. Once formed, they acted as localized sources of activation, continuously generating wavefronts and sustaining the fibrillatory activity, as illustrated in Video S1.

#### 3.4.2. Electromechanical activation, mechanical and hemodynamic response

The electromechanical, mechanical, and hemodynamic responses following the initiation of AF were compared with the healthy calibrated reference case. Electromechanical variables and pressure–volume loops are reported for the fifth simulated beat, selected as the last complete beat prior to the application of the S1–S2 protocol. Volume and flow time series are shown over three consecutive beats starting from the sixth simulated beat to capture the early arrhythmogenic dynamics following the premature stimulus.

Fig 21 presents the domain-averaged ionic and active mechanical responses. The spatially averaged intracellular calcium concentration exhibits a reduction in peak amplitude in the AF case compared with the healthy reference, with a peak decrease of 55.10%. The spatially averaged active tension shows a corresponding reduction in peak amplitude, with an approximate decrease of 74% relative to the healthy case.

**Figure 21:**
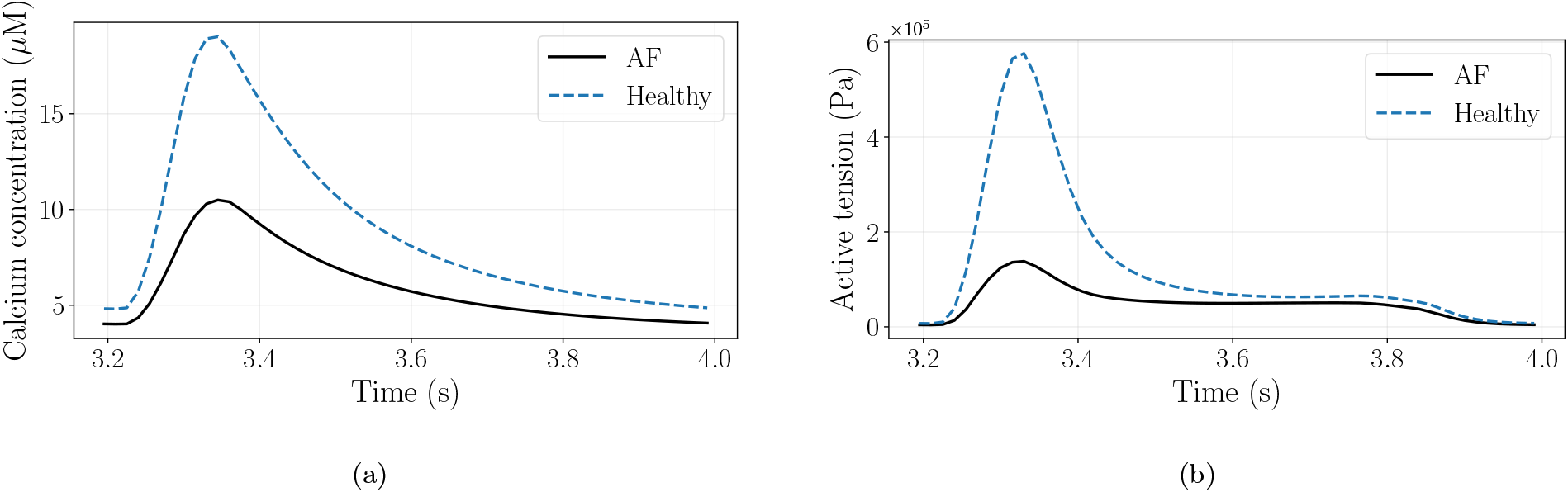
Domain-averaged electromechanical response during AF. (a) Spatially averaged intracellular calcium concentration during the fifth simulated beat. (b) Spatially averaged active tension during the fifth simulated beat. Results are shown for AF and healthy conditions.

Fig 22 summarizes the atrial mechanical response. The atrial volume–time curves show the absence of the late diastolic volume decrease associated with the atrial booster phase in the healthy condition. During the remaining phases of the cardiac cycle, atrial volumes increase and decrease following comparable patterns in both cases. This behaviour is consistently observed across the three cycles shown. The corresponding atrial pressure–volume loops reflect this difference: the healthy case displays both the A-loop and the V-loop, whereas the AF case lacks the A-loop, indicating the loss of effective atrial contraction.

**Figure 22:**
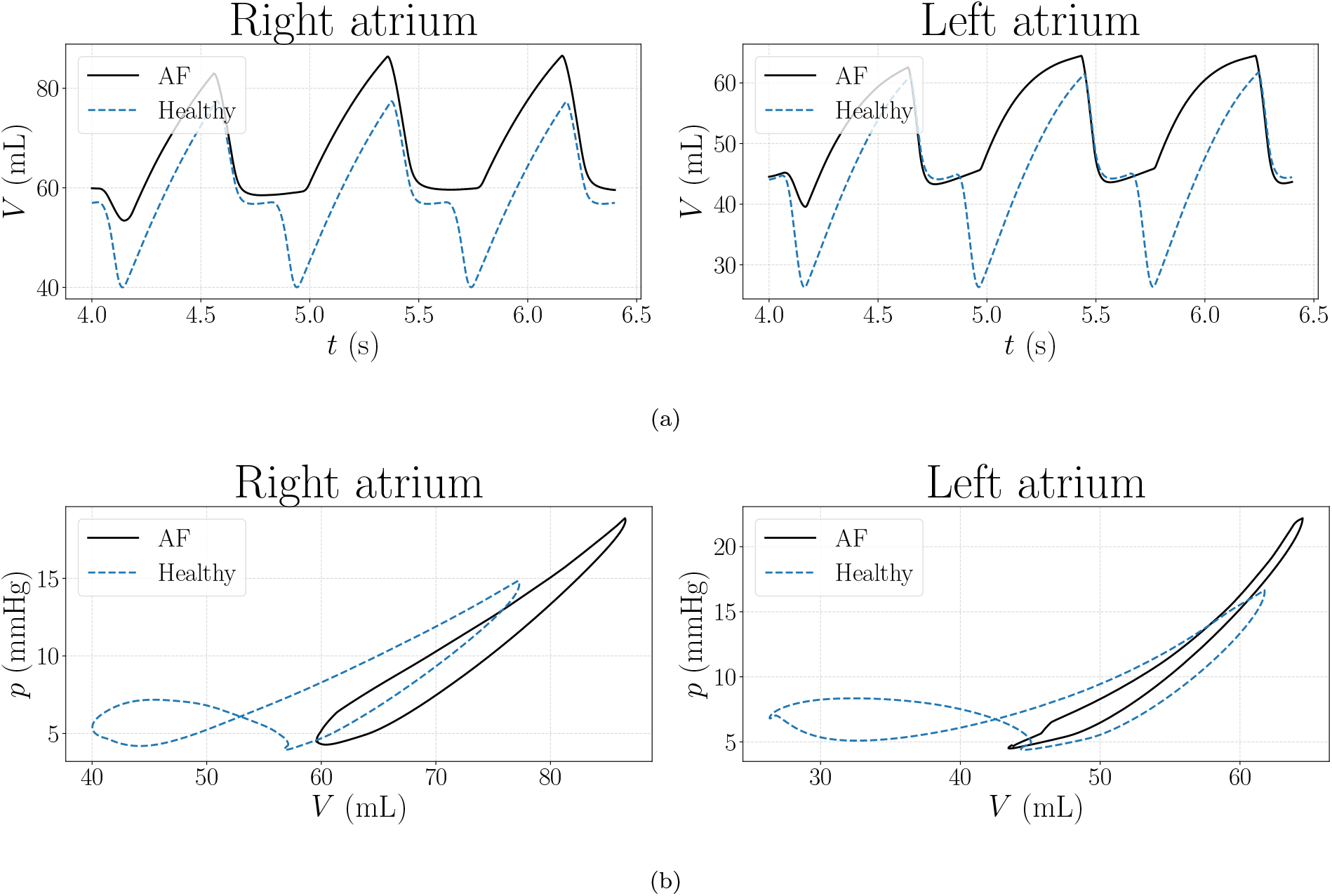
Atrial mechanical response following AF initiation. (a) Left and right atrial volumes over three consecutive simulated beats starting from the sixth beat. (b) Atrial pressure–volume loops for the fifth simulated beat. Results are shown for AF and healthy conditions.

The effects of AF are also evident in the associated hemodynamic variables accessible through the coupled 0D circulatory model, as shown in Fig 23. Atrioventricular valve flow rates show similar flux patterns between AF and healthy cases, except for the absence of the late diastolic flow component (afterwave [102]) in the AF condition. Ventricular volume–time curves indicate a reduced volume range in the AF case compared with the healthy scenario. This reduction is also reflected in the ventricular pressure–volume loops, which display an overall reduction in CO of 20.4% during the AF episode.

**Figure 23:**
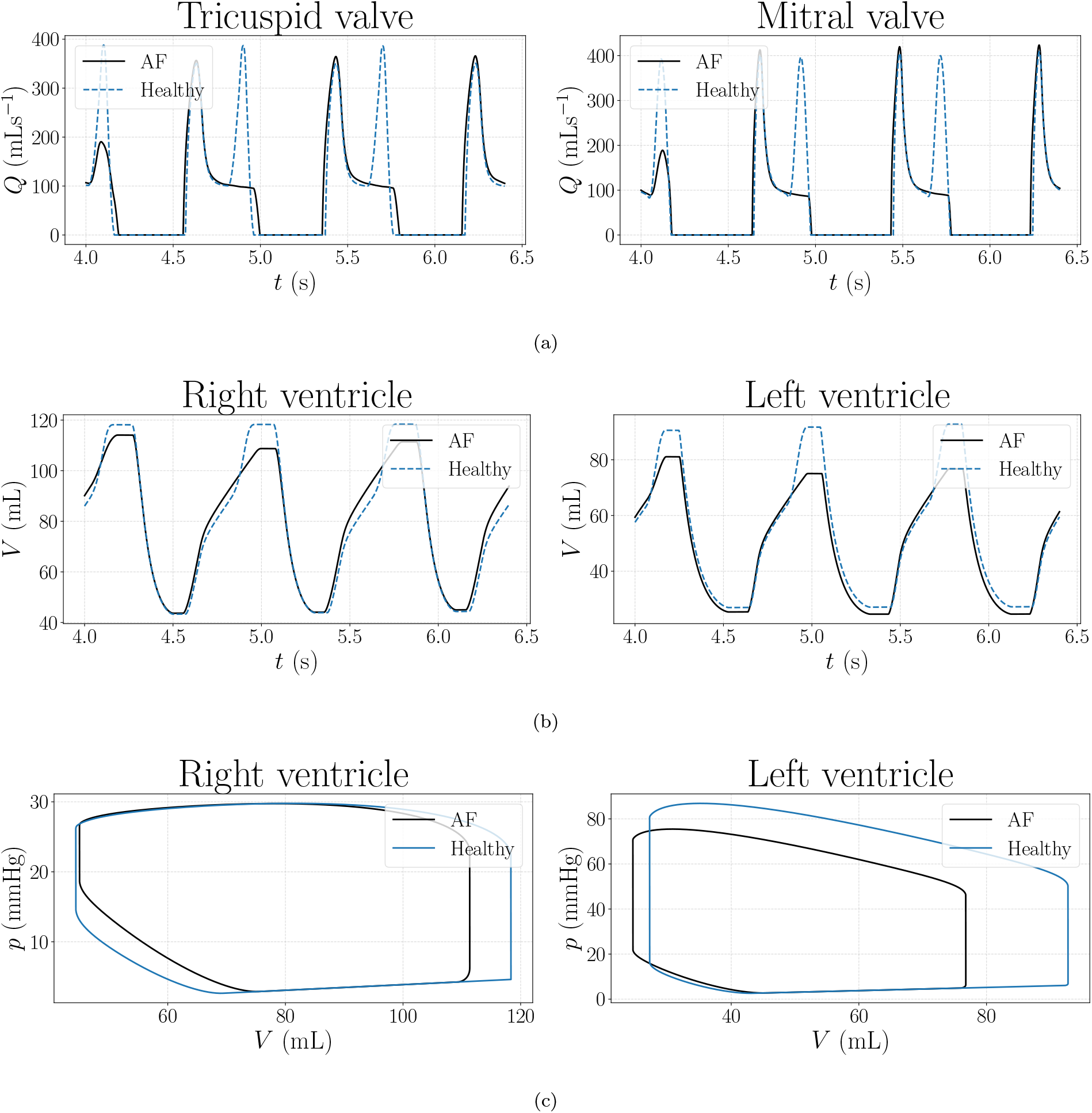
Hemodynamic and ventricular mechanical response during AF. (a) Mitral and tricuspid valve flow rates over three consecutive simulated beats starting from the sixth beat. (b) Left and right ventricular volumes over the same interval. (c) Ventricular pressure–volume loops for the fifth simulated beat. Results are shown for AF and healthy conditions.

## 4. Discussion

### 4.1. Physiological realism of atrial electromechanics

The results suggest that enforcing tight electromechanical and hemodynamic coupling within the 3D–0D biatrial framework is critical for achieving physiologically consistent atrial dynamics. By preserving the temporal hierarchy between activation, contraction, and chamber emptying, the model provides a mechanistically integrated representation of atrial function.

#### 4.1.1. Physiological atrial activation sequence

The spatiotemporal pattern of atrial electrical activation follows the physiological ordering reported by Lemery *et al*. [85]. All regional activation times during sinus rhythm fall within the corresponding experimental ranges, indicating a realistic electrical propagation sequence throughout both atria, enabled by the calibrated region-specific diffusion properties. Accurately reproducing these activation timings is essential, as they determine the spatial and temporal coordination of active tension development and therefore directly influence atrial contraction and filling dynamics.

#### 4.1.2. Atrial deformation and phase timing

The model captures a temporally organized atrial deformation pattern that allows the three functional phases to be clearly identified. The simulated atrial cycle begins with a brief contraction (booster phase), followed by a dominant reservoir phase and a shorter conduit phase, in agreement with physiological descriptions of atrial function in sinus rhythm [103].

In the presented simulation, the booster, reservoir, and conduit phases account for approximately 18%, 54%, and 29% of the cardiac cycle, respectively. These values closely match clinical measurements in healthy subjects, which typically report phase durations of approximately 20%, 45%, and 35%, respectively [104]. Importantly, these timings emerge naturally from the coupled electromechanical and hemodynamic interactions rather than being imposed explicitly, supporting the internal consistency of the model.

#### 4.1.3. Pressure–Volume loops

A key indicator of physiological realism in atrial mechanics models is the ability to reproduce characteristic atrial pressure–volume loop morphology. The model captures the figure-eight shape observed in experimental and clinical studies [105, 106, 107], with clearly distinguishable A-loops and V-loops for both atria. Reproducing this morphology remains a known challenge for atrial electromechanical models [37, 38, 108]. Only a limited number of computational studies have reported fully developed figure-eight atrial pressure–volume loops under physiological conditions [35, 34, 109]. The emergence of this morphology in the present framework reflects consistent coupling between electrical activation, tissue mechanics, and hemodynamic loading, and underscores the capacity of the model to recover integrated atrial function beyond isolated volumetric or contractile biomarkers.

#### 4.1.4. Quantitative biomarkers: Volumes and EFs

The model reproduces key quantitative features of healthy atrial function. Simulated atrial volumes and phasic EFs closely match values reported by Gao *et al*. [87]. Minimum, maximum, and pre-contraction volumes, together with reservoir, conduit, and booster EFs, all fall within physiological ranges. Importantly, this agreement is achieved simultaneously in both atria, supporting the reliability of the calibrated mechanical properties, boundary conditions, and 0D hemodynamic coupling.

### 4.2. Sensitity analysis

The sensitivity analysis offers mechanistic insight into how key parameters governing atrial contractility, passive stiffness, boundary conditions, and ventricular loading influences both atrial performance and overall cardiac function.

The reference active tension *T*_ref_ primarily modulates the atrial booster pump, as it acts as a scaling parameter for active force generation in the excitation–contraction model [65]. Variations in *T*_ref_ lead to a reduction in atrial end-systolic volume and a clear increase in booster EF, while the reservoir and conduit components of the atrial pressure–volume loop remain largely unchanged. This behaviour is physiologically consistent, since the booster phase is determined by the active atrial contraction, whereas reservoir and conduit phases are dominated by passive atrial mechanical properties rather than active force generation [110, 111].

The isotropic ground-matrix stiffness parameter *a* mainly influences atrial behaviour during the reservoir phase, reflecting its role in setting passive myocardial stiffness [78]. Increasing *a* reduces the maximum atrial volume and narrows the V-loop of the pressure–volume atrial plot, consistent with a less compliant atrium that resists passive filling. In contrast, minimum volume and booster contribution are only weakly affected, indicating that changes in intrinsic tissue stiffness tend to modulate atrial filling more markedly than active emptying. A similar trend is observed when varying the fiber stiffness parameter *a*_f_, although with slightly reduced sensitivity compared to *a*. These results suggest that, within the explored parameter range, the isotropic matrix contribution exerts a stronger influence on global atrial compliance than fiber-direction stiffness, while both components contribute to the overall deformation pattern of the chamber. This interpretation is physiologically consistent with evidence showing that increased atrial stiffness, such as that caused by fibrosis or structural remodelling, primarily impairs reservoir function while leaving active contraction phase relatively preserved [112, 113, 114].

Increasing pericardial stiffness *k*_peri_ produces a similar reduction in atrial reservoir function, with lower maximum volumes and a narrower V-loop, reflecting constrained atrial expansion during passive filling. However, unlike variations in *a*, larger values of *k*_peri_ also lead to a reduction in the A-loop area. This effect is primarily driven by a reduced atrial pressure excursion during contraction, while the end-systolic volume remains essentially unchanged. This suggests that, while high intrinsic stiffness mainly impairs passive filling, stronger pericardial constraint additionally limits pressure development during the booster phase through external mechanical loading.

An increase in active ventricular elastance *E*_act_ resulted in a more powerful ventricular contraction, reflected by an increase in ventricular EF. Under these conditions, ventricular emptying becomes more efficient, and the atria adjust their contribution through compensatory mechanisms that redistribute atrial phasic function [115, 116, 106]. In particular, the reduced area of the atrial A-loop indicates a decrease in atrial work during the booster phase, suggesting a reduced reliance on active atrial contraction to maintain ventricular filling.

In contrast, increasing ventricular passive elastance *E*_pass_ produced an opposite response. Higher passive elastance corresponds to increased ventricular stiffness, impairing ventricular filling at comparable pressures. To preserve CO, the atria compensate by augmenting their active contribution during the booster phase, as evidenced by the increase in the atrial A-loop area and the associated rise in atrial work. This behaviour is consistent with clinical observations in conditions characterized by elevated ventricular stiffness, such as restrictive cardiomyopathies [117], where atrial contraction becomes a critical determinant of ventricular filling and atrial workload is markedly increased.

### 4.3. Application to an AF scenario

A pathological episode of AF was induced within the 3D electromechanical–0D hemodynamic framework to assess how atrial electrical disorganization propagates across mechanical and hemodynamic scales, and ultimately affects global cardiac performance.

The AF initiation protocol successfully generates sustained fibrillatory activity driven by spatially heterogeneous electrical remodelling. Such heterogeneity introduces dispersion of repolarization and conduction velocity, which are well-established mechanisms underlying reentrant excitation and the maintenance of AF [118, 119, 120]. As a result, the atrial activation pattern deviates from the coherent propagation normally driven by the SAN, exhibiting marked spatial and temporal disorganization.

Beyond its impact on propagation, electrical remodelling also alters intracellular ionic dynamics. Chronic AF is known to reduce L-type calcium current leading to diminished intracellular calcium transients [91, 121, 122]. In agreement with these findings, the present simulations show a clear reduction in the spatially-averaged intracellular calcium transient under AF. Because active tension generation in the model is coupled to calcium binding dynamics, this ionic reduction translates into a marked decrease in domain-averaged active tension. This behaviour is consistent with experimental observations showing impaired atrial contractility and reduced force development in fibrillating atrial myocardium [123, 121]. Although direct quantitative comparisons remain challenging due to differences in modelling assumptions and remodelling parametrizations, the qualitative agreement supports the electromechanical consistency of the framework.

The reduction in active tension has direct mechanical consequences at the atrial level. In the healthy case, atrial volume traces exhibit two distinct emptying phases per beat: a passive reduction during the conduit phase and an additional decrease associated with active atrial contraction [110]. During AF, the second phase is absent, reflecting the loss of coordinated atrial systole. Instead, atrial volume variations are dominated by passive filling and emptying driven by venous return and ventricular suction. Consistently, atrial pressure–volume loops lose their characteristic figure-eight morphology and collapse into a single V-loop, indicating the disappearance of the booster pump contribution.

The absence of effective atrial contraction is also evident at the hemodynamic level. In the healthy case, atrioventricular valve flow profiles show both an early filling wave and a late atrial contribution [102], whereas during AF the second wave is absent [124], indicating that atrial contraction no longer contributes to ventricular filling. Consequently, ventricular enddiastolic volumes are reduced in AF, while end-systolic volumes remain nearly unchanged. This results in a reduction in stroke volume and overall CO. Clinical and experimental studies report decreases on the order of 20 to 25% in the presence of AF [125, 126, 127]. Consistently, the present simulations predict a 20.4% reduction in CO compared with the healthy case.

Compared with the electromechanical study of AF by Adeniran *et al*. [10], the present framework highlights the importance of closed-loop hemodynamics. While Adeniran *et al*. prescribed endocardial pressure, effectively suppressing conduit-driven atrial emptying, incorporation of a full 0D circulatory model in the present work allows passive atrial emptying to emerge naturally. This yields a more complete representation of how AF impairs ventricular filling and reduces overall CO.

## 5. Limitations

Some limitations of the present framework should be acknowledged.

A key limitation is the absence of mechano-electric feedback, which accounts for the influence of atrial deformation on electrophysiological behaviour. While electrical activation drives contraction through excitation-contraction coupling, the reciprocal interaction from solid mechanics to electrophysiology is neglected. Consequently, load-dependent changes in conduction and cellular ionic dynamics, which have been shown to contribute to stretch-induced electrical alterations and arrhythmogenic behaviour in atrial electromechanical and whole-heart models [37, 128, 129], are not captured.

This model uses the CRN human atrial ionic model [63] as the baseline cellular electrophysiology. Although CRN remains widely used in computational atrial studies, several updated human atrial formulations have since been proposed, incorporating revised descriptions of ionic currents and refined calcium-handling dynamics based on more recent experimental data [130, 70, 20]. Differences among ionic models can influence action potential morphology, restitution properties, calcium transient dynamics, and their response to AF remodelling, which may propagate to the electromechanical level in integrated simulations such as the one presented here. In the present study, regional heterogeneity and persistent-AF remodelling were introduced through region-specific parameter adjustments rather than inferred from patient-specific electrophysiological recordings, and therefore quantitative electrophysiological predictions may depend on the selected ionic formulation and remodelling scheme.

A limitation of this study is the use of a single biatrial anatomy for both physiological and AF simulations. Pathological behaviour is introduced through electrical remodelling and altered activation patterns, while anatomical structure is kept unchanged to isolate functional electromechanical and hemodynamic effects. As the aim of this work is not to provide subject-specific predictive validation and rather mechanistic investigation within a controlled framework, this approach is consistent with prior atrial electromechanical studies [10]. Intersubject anatomy variability is therefore not considered and remains a topic for future work.

In the present framework, atrioventricular valve rings are modelled using spring-like boundary conditions to mimic the presence of ventricles at the mechanical level, but this simplification does not capture the characteristic apical-basal excursion of the atrioventricular valve annuli or their three-dimensional deformation observed in clinical imaging studies [80, 81, 82, 83]. Alternative computational strategies, such as the use of deformable closure surfaces to seal atrioventricular openings without explicitly resolving leaflet dynamics, have been proposed in the literature [35] and can be investigated as a future direction.

Another limitation is the absence of an explicit stress-free reference configuration for the atrial myocardium. Simulations are initialized from the imaged atrial geometry, corresponding to a loaded state under physiological pressure, which is assumed as the mechanical reference. This assumption may introduce residual stresses and influence absolute strain and stress estimates, particularly in thin-walled atrial tissue [131, 132]. Recent atrial and whole-heart modelling studies have emphasized the importance of estimating a stress-free configuration through inverse or prestress procedures to ensure mechanical consistency and improve the interpretation of atrial deformation [31].

A limitation of the presented AF simulation is that ventricular activation was prescribed as regular and identical to sinus rhythm (SR) throughout the AF episode. Clinically, AF is frequently associated with an irregular ventricular response arising from atrioventricular nodal filtering of fibrillatory atrial activity [133]. Ventricular cycle length irregularity has been shown to independently affect ventricular filling and cardiac output, even at unchanged mean heart rate [134]. Incorporating AV-node–mediated ventricular activation therefore represents an important direction for future extensions of the proposed framework.

Finally, while the present framework reproduces physiologically realistic atrial pressure– volume behaviour and functional biomarkers consistent with ranges reported for healthy cohorts [85, 87], the model is neither patient-specific nor validated against subject-level data. Model parameters are chosen to represent typical atrial mechanics and hemodynamics, and model outputs are compared against population-based experimental and clinical studies. In addition, the limited availability of *in vivo* atrial motion and deformation data, currently prevents a direct quantitative validation of atrial mechanics.

## 6. Conclusions

This work introduces a multiscale biatrial 3D electromechanical–0D hemodynamic digital twin framework that enables the investigation of atrial function. The model reproduces a physiologically consistent sequence of atrial electrical activation, in agreement with experimental observations of atrial depolarization in healthy subjects. This electrical propagation pattern, coupled with mechanically consistent closed-loop hemodynamic model interaction, allows the model to capture the full atrial cycle, including reservoir, conduit, and booster pump phases.

As a result, the framework reproduces the characteristic figure-eight atrial pressure– volume loops observed experimentally and often difficult to achieve in computational studies. Model parameters are calibrated to yield atrial volumes and EFs within physiological ranges reported for a healthy cohort, supporting the ability of the framework to reproduce physiological global atrial function. Beyond matching reference conditions, a systematic sensitivity analysis is performed to distinguish the relative contributions of active contractility, passive myocardial properties, boundary conditions, and ventricular afterload, highlighting how atrial functional indices emerge from strongly coupled electromechanical and hemodynamic interactions.

Finally, the framework is applied to a pathological AF scenario by initiating an arrhythmic activation pattern and assessing its consequences at the whole-heart level. The loss of coordinated atrial activation leads to the disappearance of the booster pump phase, altered atrial pressure–volume dynamics, and impaired ventricular filling and CO, illustrating how atrial electrical dysfunction propagates across mechanical and circulatory scales. Together, these results demonstrate that the proposed framework provides a coherent and flexible methodological basis to study atrial function under physiological and pathological conditions.

## 7. Acknowledgements

The authors thankfully acknowledge the computer resources at MareNostrum V and the technical support provided by BSC (IM-2021-3-0002).

## 9. CRediT authorship contribution statement

Conceptualization: S.P.C., A.Z., V.P.G., J.S., E.C. Methodology: S.P.C., A.Z., V.P.G., E.C. Software: S.P.C., A.Z., D.L., M.V., E.C. Formal analysis: S.P.C. Investigation: S.P.C., A.Z., V.P.G., E.C. Validation: S.P.C., A.Z., D.L., E.C. Data curation: A.Z., V.P.G., J.S., M.I., E.C. Resources: V.P.G., J.S., M.I. Visualization: S.P.C., V.P.G. Supervision: A.Z., M.V., B.E., F.C., J.S., E.C. Project administration: M.V. Funding acquisition: M.V., J.S, E.C. Writing–original draft: S.P.C. Writing–review & editing: S.P.C., A.Z., V.P.G., D.L., M.V., B.E., F.C., J.S., E.C.

## 10. Ethics statement

The clinical data used to generate the patient-specific atrial mesh were provided by collaborators at Hospital Universitario y Politécnico La Fe (Valencia, Spain). The data were supplied under approval of the Ethics Committee of Hospital Universitario y Politécnico La Fe (registry number: 2023-885-1). All procedures complied with the relevant institutional and ethical guidelines for the use of clinical data in research.

## 12. Declaration of generative AI and AI-assisted technologies in the manuscript preparation process

During the preparation of this work the authors used ChatGPT in order to assist with language editing, minor text restructuring, and formatting suggestions. After using this tool, the authors reviewed and edited the content as needed and take full responsibility for the content of the published article.

## 13. Supplementary materials

Supplementary material associated with this article can be found in the online version.

## Footnotes

1 https://seg3d.readthedocs.io/en/latest/

2 https://www.paraview.org/

3 https://www.blender.org/

4 https://github.com/PyMesh/quartet

5 https://www.beta-cae.com/ansa.htm

6 https://www.bsc.es/marenostrum/marenostrum-5

