## Supplementary material for "A biatrial digital twin integrating electrophysiology, mechanics, and circulation: from physiology to atrial fibrillation": S1 Table

**S1 Table. Passive mechanics parameters of the Holzapfel–Ogden law.**

| Parameter | Symbol | Unit | Source | Reference value |
|---|---|---|---|---|
| Isotropic ground-matrix stiffness | $a$ | Pa | [1] | 650 |
| Isotropic ground-matrix nonlinearity | $b$ | – | [1] | 5 |
| Fiber stiffness | $a_{\mathrm{f}}$ | Pa | [1] | 3260 |
| Fiber nonlinearity | $b_{\mathrm{f}}$ | – | [1] | 5 |
