## Supplementary material for "A biatrial digital twin integrating electrophysiology, mechanics, and circulation: from physiology to atrial fibrillation": S2 Table

**S2 Table. Active mechanics parameters of the Land–Niederer model.**

| Parameter | Symbol | Unit | Source | Reference value |
|---|---|---|---|---|
| Calcium unbinding rate | $k_{\mathrm{TRPN}}$ | $\mathrm{ms}^{-1}$ | [1] | 0.1 |
| Cooperativity ($\mathrm{Ca}^{2+}$–TnC) | $n_{\mathrm{TRPN}}$ | | [1] | 2 |
| Half-activation Ca level | $[\mathrm{Ca}]_{T50}^{\mathrm{ref}}$ | $\mu$M | [2] | 0.86 |
| Tropomyosin unblocking rate | $k_u$ | $\mathrm{ms}^{-1}$ | [1] | 1 |
| Cooperativity (Tm) | $n_{\mathrm{Tm}}$ | | [1] | 5 |
| Weak-to-strong XB rate | $k_{ws}$ | $\mathrm{ms}^{-1}$ | [2] | 0.036 |
| Unbound-to-weak XB rate | $k_{uw}$ | $\mathrm{ms}^{-1}$ | [2] | 0.546 |
| Weak-state fraction | $r_w$ | | [1] | 0.5 |
| Strong-state fraction | $r_s$ | | [1] | 0.25 |
| Strong XB unbinding slope | $\gamma_s$ | | [1] | 0.0085 |
| Weak XB unbinding slope | $\gamma_w$ | | [1] | 0.615 |
| Distortion decay factor | $\phi$ | | [1] | 2.23 |
| Instantaneous stiffness scale | $A_{\mathrm{eff}}$ | | [1] | 25 |
| Length-dependent max tension | $\beta_0$ | | [1] | 2.3 |
| Length-dependent Ca sensitivity | $\beta_1$ | | [1] | -2.4 |
| Reference tension | $T_{\mathrm{ref}}$ | kPa | [1] | 120 |
