## Supplementary material for "A biatrial digital twin integrating electrophysiology, mechanics, and circulation: from physiology to atrial fibrillation": S3 Table

**S3 Table. Parameters of the 0D closed-loop circulation model.**

| **Parameter** | **Symbol** | **Unit** | **Source** | **Reference value** |
|---|---|---|---|---|
| Chamber elastances | | | | |
| LV active elastance | $E_{\mathrm{LV,act}}$ | mmHg/mL | [1] | 2.75 |
| LV passive elastance | $E_{\mathrm{LV,pass}}$ | mmHg/mL | [1] | 0.08 |
| RV active elastance | $E_{\mathrm{RV,act}}$ | mmHg/mL | [1] | 0.55 |
| RV passive elastance | $E_{\mathrm{RV,pass}}$ | mmHg/mL | [1] | 0.05 |
| LV contraction onset | $t_{\mathrm{LV,c}}$ | s | [1] | 0.00 |
| RV contraction onset | $t_{\mathrm{RV,c}}$ | s | [1] | 0.00 |
| LV contraction duration | $d_{\mathrm{LV,c}}$ | s | [1] | 0.34 |
| RV contraction duration | $d_{\mathrm{RV,c}}$ | s | [1] | 0.34 |
| LV relaxation duration | $d_{\mathrm{LV,r}}$ | s | [1] | 0.15 |
| RV relaxation duration | $d_{\mathrm{RV,r}}$ | s | [1] | 0.15 |
| Vascular compartments | | | | |
| Systemic arterial compliance | $C_{\mathrm{SYS,AR}}$ | mL/mmHg | [2] | 1.2 |
| Systemic venous compliance | $C_{\mathrm{SYS,VEN}}$ | mL/mmHg | [2] | 60.0 |
| Pulmonary arterial compliance | $C_{\mathrm{PUL,AR}}$ | mL/mmHg | [2] | 10.0 |
| Pulmonary venous compliance | $C_{\mathrm{PUL,VEN}}$ | mL/mmHg | [2] | 16.0 |
| Systemic arterial inductance | $L_{\mathrm{SYS,AR}}$ | mmHg·s$^2$/mL | [2] | $5 \times 10^{-3}$ |
| Systemic venous inductance | $L_{\mathrm{SYS,VEN}}$ | mmHg·s$^2$/mL | [2] | $5 \times 10^{-4}$ |
| Pulmonary arterial inductance | $L_{\mathrm{PUL,AR}}$ | mmHg·s$^2$/mL | [2] | $5 \times 10^{-4}$ |
| Pulmonary venous inductance | $L_{\mathrm{PUL,VEN}}$ | mmHg·s$^2$/mL | [2] | $5 \times 10^{-4}$ |
| Systemic arterial resistance | $R_{\mathrm{SYS,AR}}$ | mmHg·s/mL | [2] | 0.8000 |
| Systemic venous resistance | $R_{\mathrm{SYS,VEN}}$ | mmHg·s/mL | [2] | 0.2600 |
| Pulmonary arterial resistance | $R_{\mathrm{PUL,AR}}$ | mmHg·s/mL | [2] | 0.1625 |
| Pulmonary venous resistance | $R_{\mathrm{PUL,VEN}}$ | mmHg·s/mL | [2] | 0.1625 |
| Valves | | | | |
| Minimum valve resistance | $R_{\mathrm{min}}$ | mmHg·s/mL | [2] | 0.0075 |
| Maximum valve resistance | $R_{\mathrm{max}}$ | mmHg·s/mL | [2] | 75000 |
