## Supplementary figures and images for "A biatrial digital twin integrating electrophysiology, mechanics, and circulation: from physiology to atrial fibrillation"

### S1 Video

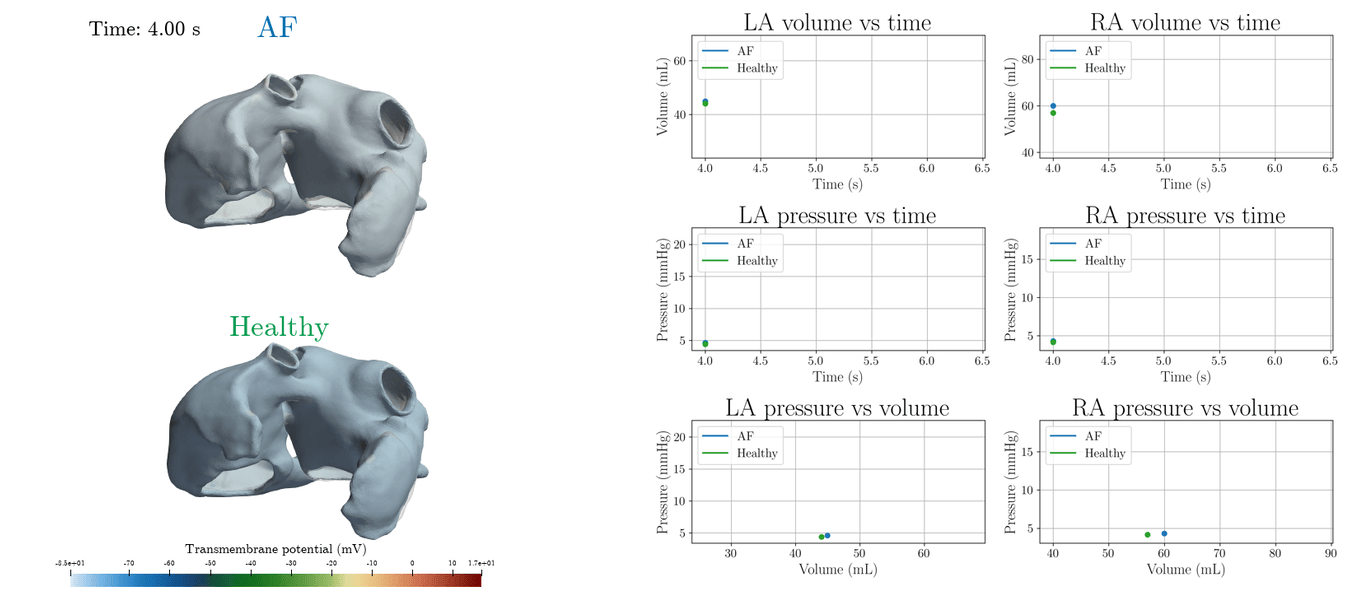
